# Spatial reasoning via recurrent neural dynamics in mouse retrosplenial cortex

**DOI:** 10.1101/2022.04.12.488024

**Authors:** Jakob Voigts, Ingmar Kanitscheider, Nicholas J. Miller, Enrique H.S. Toloza, Jonathan P. Newman, Ila R. Fiete, Mark T. Harnett

## Abstract

From visual perception to language, sensory stimuli change their meaning depending on prior experience. Recurrent neural dynamics can interpret stimuli based on externally cued context, but it is unknown whether similar dynamics can compute and employ internal hypotheses to resolve ambiguities. Here, we show that mouse retrosplenial cortex (RSC) can form hypotheses over time and perform spatial reasoning through recurrent dynamics. In our task, mice navigated using ambiguous landmarks that are identified through their mutual spatial relationship, requiring sequential refinement of hypotheses. Neurons in RSC and in artificial neural networks encoded mixtures of hypotheses, location, and sensory information, and were constrained by robust low dimensional dynamics. RSC encoded hypotheses as locations in activity space with divergent trajectories for identical sensory inputs, enabling their correct interpretation. Our results indicate that interactions between internal hypotheses and external sensory data in recurrent circuits can provide a substrate for complex sequential cognitive reasoning.

## Introduction

External context can change the processing of stimuli^1,2^ via recurrent neural dynamics^3^. To study how hypotheses can serve as internal context signals, we developed a task that requires sequential integration of ambiguous stimuli across time and space^4^. Freely moving mice have to distinguish between two perceptually identical landmarks, formed by identical dots on a computer-display arena floor, by sequentially visiting them and reasoning about their relative locations. The landmarks were separated by <180 degrees in an otherwise featureless circular arena (50 cm diameter), to create a clockwise (‘a’) and a counterclockwise (‘b’) landmark. Across trials, the relative angle between landmarks was fixed and the same relative port was always the rewarded one; within trials, the locations of landmarks was fixed. The mouse’s task was to find and nose-poke at the ‘b’ landmark for water reward (‘b’ was near one of 16 identical reward ports spaced uniformly around the arena; other ports caused a time-out). At most one landmark was visible at a time (enforced by tracking mouse position and modulating landmark visibility based on relative distance, see Methods, Supplementary Fig. 1). Each trial began with the mouse in the center of the arena in the dark (‘LM0’ phase, Fig. 1b), without knowledge of its initial pose. In the interval after first encountering a landmark (‘LM1’ phase), an ideal agent’s location uncertainty is reduced to two possibilities, but there is no way to disambiguate whether it saw ‘a’ or ‘b’. After seeing the second landmark, an ideal agent could infer landmark identity (‘a’ or ‘b’; this is the ‘LM2’ phase, Fig. 1b) by estimating the distance and direction traveled since the first landmark and comparing those with the learned relative layout of the two landmarks; thus, an ideal agent can use sequential spatial reasoning to localize itself unambiguously. To randomize the absolute angle of the arena at the start of each new trial (and thus avoid use of any olfactory or other cues), mice had to complete a separate instructed visually-guided dot-hunting task, after which the landmarks were randomly rotated together (Supplementary Fig. 1). Mice learned the task (Fig.1c, p < 0.0001 on all mice, Binomial test vs. random guessing), showing that they learn to form hypotheses about their position during the LM1 phase, retain and update these hypotheses with self-motion information until they encounter the second (perceptually identical) landmark, and use them to disambiguate location and determine the rewarded port.

**Figure 1.**
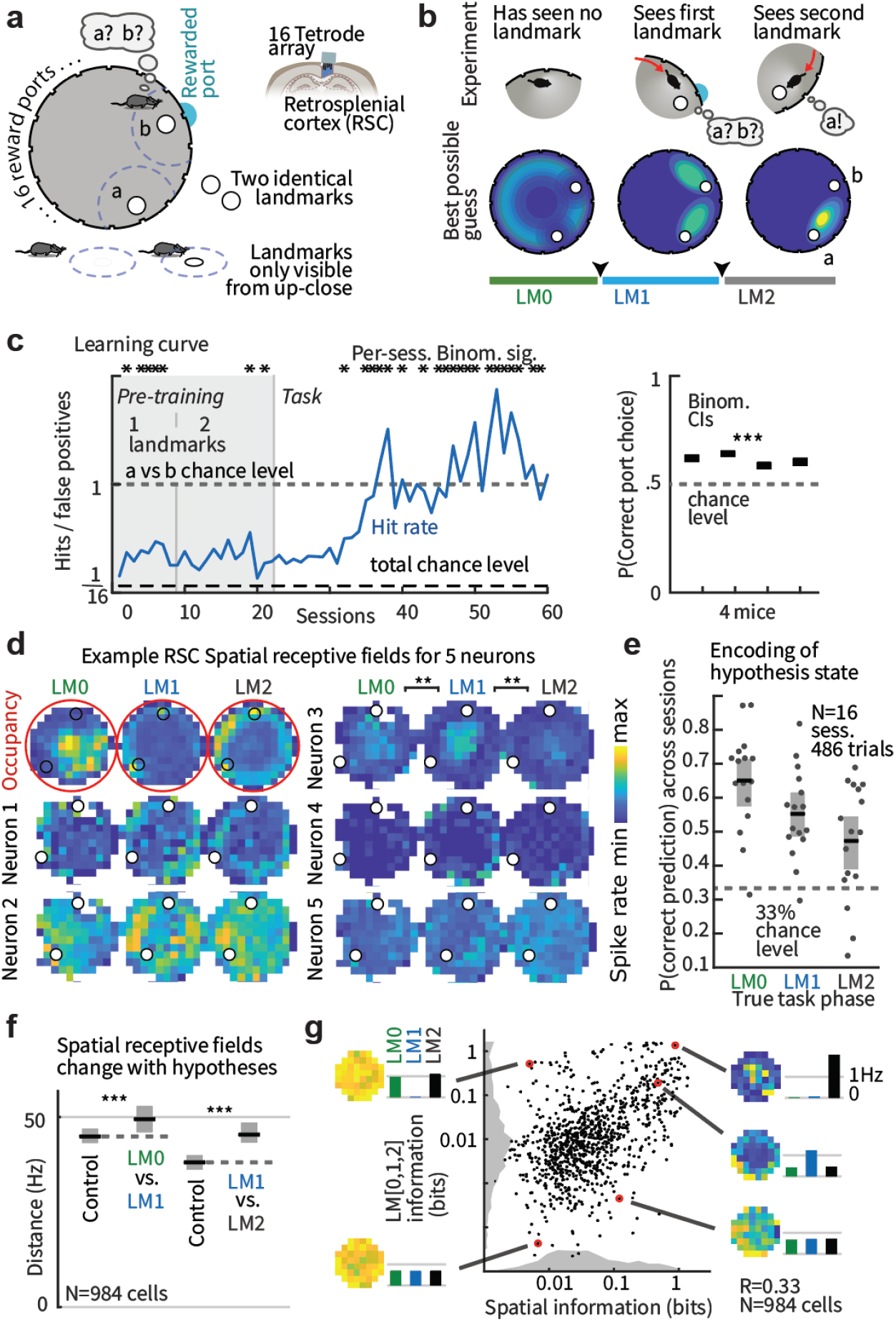
Retrosplenial cortex represents spatial information conjunctively with hypothesis states during navigation with locally ambiguous landmarks. (**a**) Two perceptually identical landmarks are only visible from close-up and their identity is only defined by their relative location. One of 16 ports, at landmark ‘b’, delivers reward in response to a nose-poke; the animal must infer which of the two landmarks is ‘b’ to receive reward; wrong pokes result in timeout. Tetrode array recordings in retrosplenial cortex (RSC) yield 50-90 simultaneous neurons. (**b**) Schematic example trial (top) and best possible guesses of the mouse position (bottom). ‘LM0,1,2’ denotes task phases when the mouse has seen 0, 1, or 2 landmarks and could infer its position with decreasing uncertainty. (**c**) Left: Example training curve showing P_hit_/P_false-positive_; random chance level is 1/16 for 16 ports. Mice learned the task at values > 1, showing they could disambiguate between the two sequentially visible landmarks. This requires the formation, maintenance, and use of spatial hypotheses. Summary statistics at right show binomial CIs on last half of sessions for all 4 mice. (**d**) Mouse location heatmap from one session (red) with corresponding spatial firing rate profiles for 5 example cells. (**e**) Task phase (corresponding to hypothesis states, i.e. panel b), can be decoded from RSC firing rates. Shading: 95% confidence intervals (CIs). (**f**) Spatial coding changes between LM1 and LM2 phases (Euclidean distances between spatial firing rate maps, control within vs. across condition, Supplementary Fig.5a for test via decoding). (**g**) Spatial vs. task phase information content of all neurons and encoding for example cells. Grey: sum-normalized histograms.

We hypothesized that RSC, which integrates self-motion^5^, position^6–8^, and sensory^9^ inputs, could perform this computation. RSC is causally required to process landmark information^10^, and we verified that RSC is required for integrating spatial hypotheses with visual information but not for direct visual search with no memory component (Supplementary Fig. 2).

### Spatial hypotheses are encoded conjunctively with other navigation variables in RSC

We recorded 50-90 simultaneous neurons in RSC during navigational task performance using tetrode array drives^11^ (Fig. 1a) and behavioral tracking (see Methods, Supplementary Figs. 1,3,4). RSC neurons encoded information about both the mouse’s location (Fig. 1d) and about the task phase, corresponding to possible location hypotheses (Fig. 1d,e). This hypothesis encoding was not restricted to a separate population: most cells encoded both hypothesis state as well as the animal’s location (Fig. 1g). This encoding was distinct from the encoding of landmark encounters in the interleaved dot-hunting task and was correlated per-session with behavioral performance (Supplementary Fig. 6). The encoding of mouse location changed across task phases (Fig. 1d,f), similar to the conjunctive coding for other spatial and task variables in RSC^6^. This mixed encoding suggests that RSC can transform new ambiguous sensory information into unambiguous spatial information through the maintenance and contextual use of internally generated spatial hypotheses.

### Hypothesis-dependent spatial computation via recurrent dynamics

To test whether recurrent neural networks can solve sequential spatial reasoning tasks that require hypothesis formation, and to provide insight into how this might be achieved in the brain, we trained a recurrent artificial neural network (ANN) on a simplified 1-dimensional version of the task, since the relevant position variable for the landmarks was their angular position (Fig. 2b; inputs were random noisy velocity trajectories and landmark positions, but not their identity). The ANN performed as well as a near Bayes-optimal particle filter (Fig. 2b), outperforming path integration with correction (corresponding to continuous path integration^12,13^ with boundary/landmark resetting^14,15^), and represented multi-modal hypotheses, transitioning from a no-information state (in LM0) to a bimodal two-hypothesis coding state (LM1), and finally to a full information, one-hypothesis coding state (LM2) (Fig. 2c,d, Supplementary Fig.7). This result shows that recurrent neural dynamics are sufficient to internally generate, retain, and apply hypotheses to reason across time based on ambiguous sensory and motor information, with no external context inputs.

**Figure 2.**
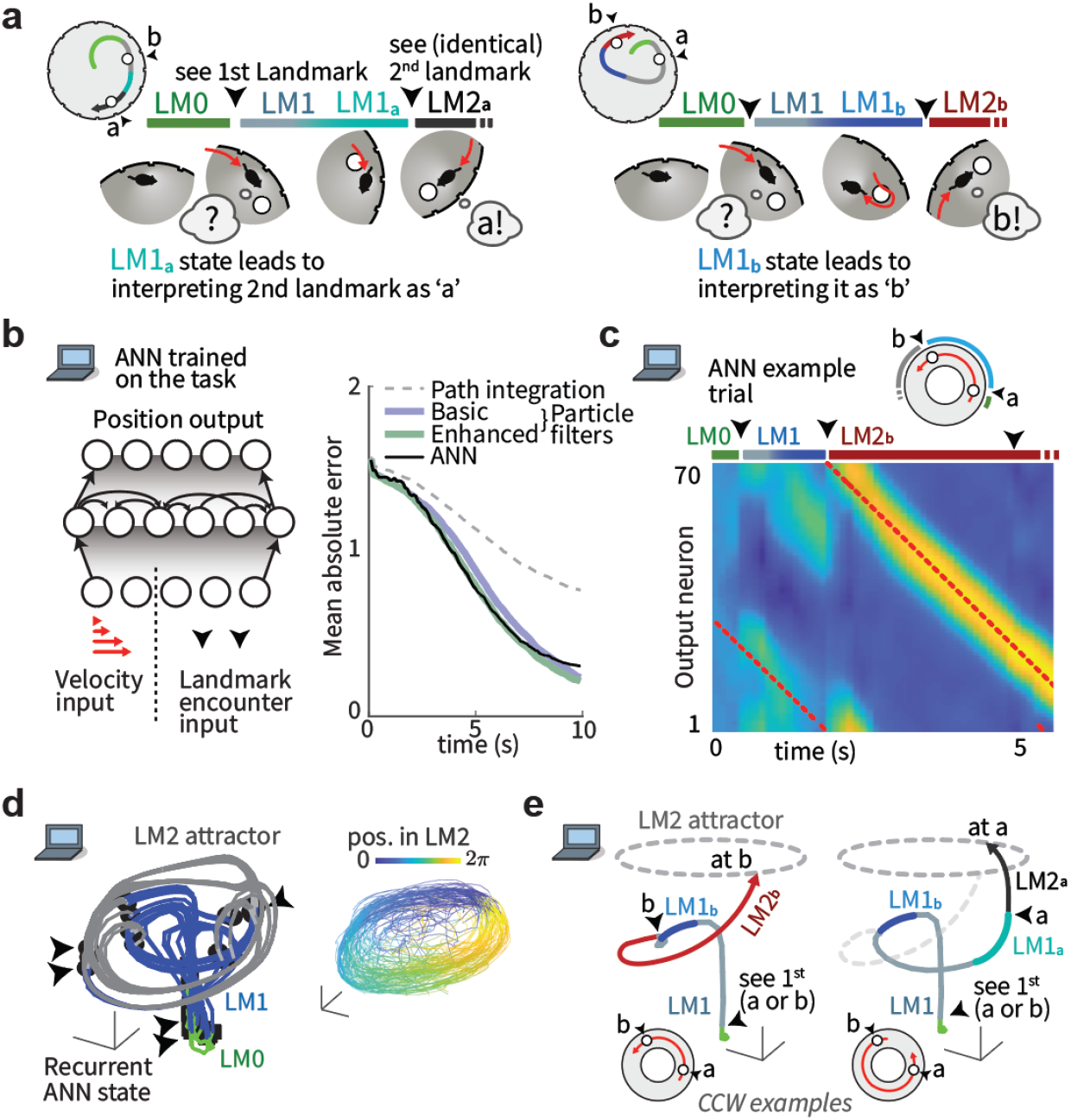
Recurrent neural dynamics can be used to navigate via locally ambiguous landmarks by forming and employing multimodal hypotheses. (**a**) Schematic examples of hypothesis-dependent landmark interpretation. Left: Mouse encounters first landmark, then identifies the second as ‘a’ based on the short relative distance. Right: A different path during LM1 leads the mouse to a different hypothesis state, and to identify the perceptually identical second landmark as ‘b’. Hypothesis states preceding LM2 are denoted LM1_a_ and LM1_b_, depending on the identity of the second landmark. (**b**) Structure of an ANN trained on the task. Inputs encode velocity and landmarks. Right: Mean absolute localization error averaged across test trials, for random trajectories. (**c**) Activity of output neurons ordered by preferred location shows transition between LM0,1,2 phases. Red: true location. During LM1 (when the agent has only seen one landmark), two hypotheses are maintained, with convergence to a stable unimodal location estimate in LM2 after encountering the second landmark. (**d**) 3D projection of ANN hidden neuron activities. During LM2, angular position in neural state space reflects position estimate encoding. (**e**) Example ANN trajectories for two trials show how identical visual input (black arrows) leads the activity to travel to different locations on the LM2 attractor because of different preceding LM1_a/b_ states.

Instantaneous position uncertainty (variance derived from particle filter) could be decoded from ANN activity (Supplementary Fig. 8a), analogous to RSC (Fig.1e). Both ANN and RSC neurons encoded multiple navigation variables conjunctively (Supplementary Fig.5b), and preferentially represented landmark locations (Supplementary Fig.5c; consistent with overrepresentation of reward sites in hippocampus^16,17^), and transitioned from encoding egocentric landmark-relative position during LM1 to a more allocentric encoding during LM2 (Supplementary Fig. 9). Average spatial tuning curves of ANN neurons were shallower in the LM1 state relative to LM2, corresponding to trial-by-trial ‘disagreements’ between neurons, evident as bimodal rates per location. RSC rates similarly became less variable across trials per-location in LM2 (Supplementary Fig. 10), indicating that in addition to the explicit encoding of hypotheses/uncertainty (Fig.1e,g), there is a higher degree of trial-to-trial variability in RSC as a function of spatial uncertainty.

The ANN computed, retained, and used multi-modal hypotheses to interpret otherwise ambiguous inputs: after encountering the first landmark, the travel direction and distance to the second is sufficient to identify it as ‘a’ or ‘b’ (Figs. 1b, 2a). There are four possible scenarios for the sequence of landmark encounters: ‘a’ then ‘b’, or ‘b’ then ‘a’, for CW or CCW travel directions respectively. To understand the mechanism by which hypothesis encoding enabled disambiguation, we examined the moment when the second landmark becomes visible and can be identified (Fig. 2a). We designate LM1 states in which the following second landmark is ‘a’ as ‘LM1_a_’ and those that lead to ‘b’ as ‘LM1_b_’. Despite trial-to-trial variance resulting from random exploration trajectories and initial poses, ANN hidden unit activity fell on a low-dimensional manifold (correlation dimension d ≈ 3, Fig. 3d) and could be well captured in a 3D embedding via PCA (Fig. 2d). Activity states during the LM0,1,2 phases (green, blue, grey, respectively) were distinct, and transitions between phases (mediated by identical LM encounters; black arrows) clustered into discrete locations. Examining representative trajectories (for the CCW case, Fig.2e) reveals that LM1_a_ and LM1_b_ states are well-separated in activity space. If the second landmark appears at the shorter CCW displacement (corresponding to the ‘a’ to ‘b’ interval), the state jumps to the ‘b’ coding point on the LM2 attractor (Fig.2e). On the other hand, the absence of a landmark at the shorter displacement causes the activity to traverse LM1_a_, until the 2^nd^ landmark causes a jump onto the ‘a’ coding location on the LM2 attractor. In both cases, an identical transient landmark input pushes the activity from distinct hypothesis-encoding regions of activity space onto different appropriate locations in the LM2 state, constituting successful localization.

We next consider the nature of the dynamics and representation that allows the circuit to encode the same angular position variables across LM1 and LM2 regimes while also encoding the different hypotheses required to disambiguate identical landmarks. Does the latter drive the network to functionally reorganize throughout the computation? Or, does the former, together with the need to maintain and use the internal hypotheses across time, require the network to exhibit stable low-dimensional recurrent attractor dynamics? To test this, we computed the pairwise correlations of the ANN activity states (Fig. 3a) and found them to be well conserved across LM1 and LM2 states. As these correlation matrices are the basis for projections into low-dimensional space, this shows that the same low-dimensional dynamics were maintained, despite spanning different computational and hypothesis-encoding regimes (metastable two-state encoding with path integration in LM1 vs. stable single-state path integration unchanged by further landmark inputs in LM2, Supplementary Fig. 7). Low-dimensional pairwise structure was also conserved across different landmark configurations and varied ANN architectures, and the low-dimensionality of ANN states was robust to large perturbations (Supplementary Figs.12, 15). In sum, these computations were determined by one stable set of underlying recurrent network dynamics, which, together with appropriate self-motion and landmark inputs, can maintain and update hypotheses to disambiguate identical landmarks over time, with no need for external context inputs.

**Figure 3.**
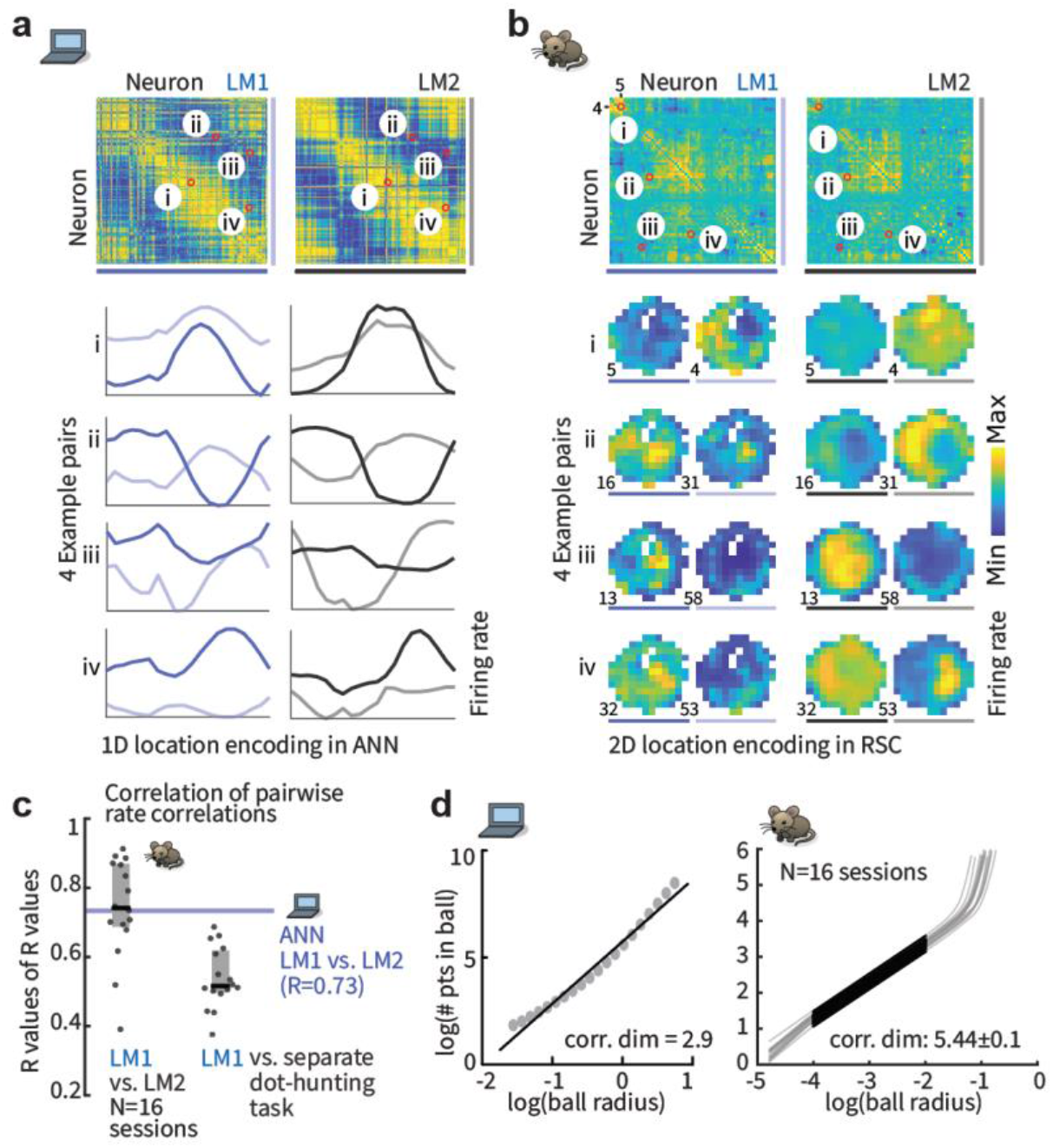
Stable low-dimensional dynamics for hypothesis-based stimulus disambiguation. (**a**) Correlation structure in ANN activity is maintained across task phases, indicating maintained low-dimensional neural dynamics across different computational regimes. Top: Pairwise ANN tuning correlations in LM1 and LM2 (same ordering, by preferred location). Bottom: tuning curve pairs (normalized amplitude). (**b**) Same analysis as a, but for RSC in one session (N = 64 neurons, computed on entire spike trains). The re-organization of spatial coding as hypotheses are updated (See also Fig. 1d,f) is constrained by the stable pairwise structure of RSC activity. Neurons remain correlated (1^st^ and 2^nd^ pair), or anti-correlated (3^rd^ and 4^th^ pair), across LM1 and LM2. (**c**) Summary statistics (sessions and quartiles) for maintenance of correlations across task phases. This also extends to a separate visually guided dot-hunting task (see also Supplementary Fig. 13). (**d**) Activity in both the ANN and RSC is locally low-dimensional (via correlation dimension, Supplementary Fig.13 for analysis via PCA).

### RSC fulfills requirements for hypothesis-dependent spatial computation via recurrent dynamics

We hypothesized that RSC and its reciprocally connected brain regions may, similarly to the ANN, use internal hypotheses to resolve landmark ambiguities via recurrent dynamics. To test this, we first computed pairwise rate correlations and found a preserved structure between LM1 and LM2, as in the ANN (median R or Rs = 0.74 in RSC, vs 0.73 in ANN, Fig. 3c). Firing rates could be well-predicted from rates of other neurons, using pairwise rate relationships across task phases; this maintained structure also extended to the visual dot-hunting behavior (Supplementary Fig. 13), indicating that RSC activity is coordinated by the constraints of stable recurrent neural dynamics and not a feature of a specific behavioral task.

Consistent with highly conserved cell-cell relationships, RSC population activity was low-dimensional (∼6 significant principal components, and correlation dim. ∼5.4, Fig. 3d, Supplementary Fig. 13), similar to findings in hippocampus^18^. Together, we find that despite significant changes in neural encoding as different hypotheses are entertained across task phases (Fig.1d,f, Supplementary Fig. 5a) and across different tasks (Supplementary Fig. 6), the evolution of firing rates in RSC is constrained by stable attractor dynamics that could implement qualitatively similar computations as the ANN (Fig. 4a).

**Figure 4.**
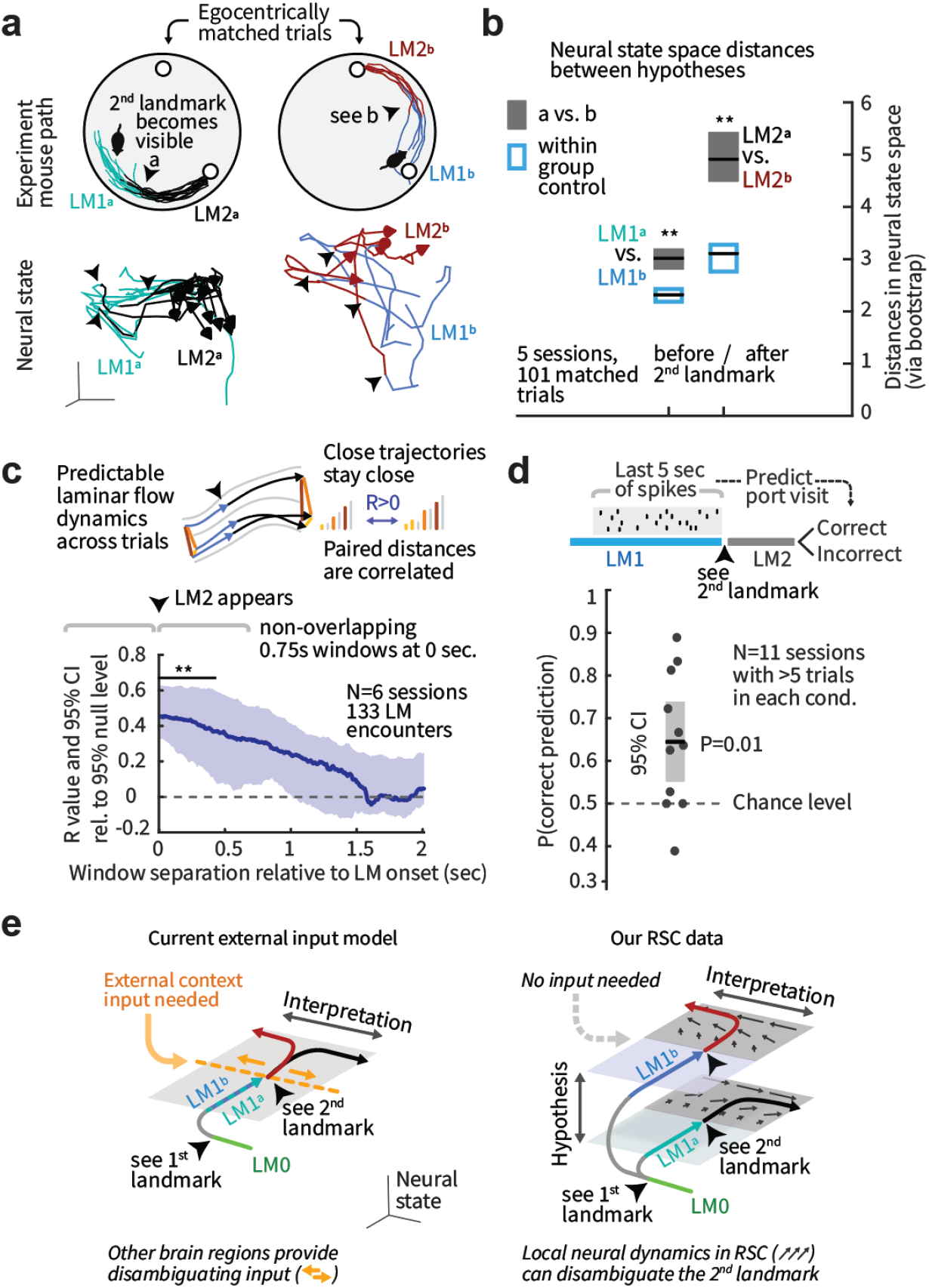
RSC exhibits stable attractor dynamics sufficient for computing hypothesis-dependent landmark identity. (**a**) Top: To study the encoding of hypotheses and its impact on sensory processing without sensory or motor confounds, we used trials with matched egocentric paths just before and after the second landmark (‘a’ or ‘b’) encounter. One example session is shown. Bottom: corresponding 3-D neural state space trajectories (via isomap). RSC latent states do not correspond directly to those of the ANN. (**b**) RSC represents the difference between LM1_a_ and LM1_b_, and between subsequent LM2 states, as in the ANN (Fig. 2e, Supplementary Figs. 7-9). Blue: within-group and red: across-group distances in neural state space. CIs via bootstrap. State can also be decoded from raw spike rates (Supplementary Fig. 14j). (**c**) Neural dynamics in RSC are smooth across trials: pairwise distances between per-trial spike counts in a 750ms window before LM2 is visible remain correlated with later windows. (**d**) RSC activity preceding the second landmark encounter predicts correct/incorrect port choice (cross-validated regression trees). (**e**) Schematic of potential computational mechanisms. Left: If RSC encoded only current spatial and sensorimotor states and no hypotheses (LM1_a_ or LM2_b_, derived from seeing the first landmark and self-motion integration that lead to identifying the second landmark as ‘a’ or ‘b’), an external disambiguating input is needed. Right: Because hypotheses are encoded, and activity follows stable attractor dynamics (Fig. 3), ambiguous visual inputs can drive the neural activity to different positions, disambiguating landmark identity in RSC analogously to the ANN.

We next examined the dynamical evolution of neural states in RSC during the spatial reasoning process. States evolved at speeds correlated with animal locomotion (Supplementary Fig. 14a), consistent with the observation that hypotheses are updated by self-motion in between landmark encounters (Fig. 1) and were driven by landmarks (Supplementary Fig. 14a), consistent with findings in head-fixed tasks^10^. To test the hypothesis that different neural states in LM1_a_ and LM1_b_ together with stable low-dimensional attractor dynamics can resolve the identity of the second perceptually ambiguous landmark, we identified subsets of trials in which mouse motion around the LM1 to LM2 transition was closely matched and aligned them in time to the point when the second landmark became visible (Fig. 4a, Supplementary Fig. 14). In these trials, locomotion and visual inputs are matched, but the preceding hypothesis state (LM1_a_ or _b_) differs. RSC firing rates differed between LM1_a_ and LM1_b_ states, as did subsequent rates in LM2 (Fig. 4b, comparing within-to across-group distances in neural state space across matched trials, and by decoding state from firing rates: Supplementary Fig. 14). The evolution of RSC firing rates was also predictable across trials such that neighboring trials remained nearby in activity space (Fig. 4c), which further confirms stable recurrent dynamics and indicates a topological organization of abstract task variables^18^. This indicates that stably maintained hypothesis-encoding differences in firing over LM1 in the low-dimensional attractor dynamics could interact with identical visual landmark inputs to push neural activity from distinct starting points in neural state space to points that correspond to correct landmark interpretations, as in the ANN.

Further, we observed that neural trajectories from LM1_a_ that were close in activity-space to LM1_b_ were dragged along LM1_b_ trajectories and vice-versa (they had similar movement directions, Supplementary Fig. 14g,h), suggesting that behavioral landmark identification outcomes might be affected by how hypotheses were encoded in RSC during LM1. We tested this hypothesis and found that RSC activity in LM1 (last 5 sec preceding LM2) was indeed predictive of the animal’s behavioral choice of the correct vs. incorrect port (Fig. 4d). Notably, this behaviorally predictive hypothesis encoding was absent during training in sessions with low task performance (Supplementary Fig. 6), indicating that the dynamical structures and hypothesis states observed in RSC were task-specific and acquired during learning. Our unrestrained non-stereotyped behavior is not amenable to direct comparison of activity trajectories in ANNs and the brain as others have done in highly stereotyped trials of macaque behavior^3^. Instead, we found that the dynamics of firing rates in mouse RSC are consistent with, and sufficient for, implementing hypothesis-based disambiguation of identical landmarks using a similar computational mechanism as observed in the ANN.

## Discussion

We report that RSC represents internal spatial hypotheses, sensory inputs, and their interpretation and fulfills the requirements for computing and using hypotheses to disambiguate landmark identity via local recurrent dynamics. Specifically, we found that low-dimensional recurrent dynamics were sufficient to perform spatial reasoning (i.e. to form, maintain, and use hypotheses to disambiguate landmarks over time) in an ANN (Fig. 2). We then found that RSC fulfills the requirements for such dynamics, i.e. encoding of the required variables (Figs. 1, 4) with stable low-dimensional (Fig. 3) and smooth dynamics that predicted behavioral outcomes (Fig. 4).

We observed that local dynamics in RSC can disambiguate sensory inputs based on internally generated and maintained hypotheses without relying on external context inputs at the time of disambiguation (Fig. 4), indicating that RSC can derive hypotheses over time and combine these hypotheses with accumulating evidence from the integration of self-motion (e.g., paths after the first landmark encounter) and sensory stimuli to solve a spatiotemporally extended spatial reasoning task. These results do not argue for RSC as an exclusive locus of such computations. There is evidence for parallel computations, likely at different levels of abstraction, across sub-cortical^19^ and cortical regions such as PFC^1,20,21^, PPC^22^, LIP^23^, and visual^24,25^ areas. Further, hippocampal circuits contribute to spatial computations beyond representing space by learning environmental topology^26^ and constraining spatial coding via attractor dynamics^27,28,18^ shaped by prior experience^29^. Finally, the landmark disambiguation that we observed likely interacts with lower sensory areas^30^ and action selection computations^31,32^.

The emergence of conjunctive encoding, explicit hypothesis codes, and similar roles for dynamics across RSC and the ANN suggests that spatial computations and, by extension, cognitive processing in neocortex may be constrained by simple cost functions^33^, similar to sensory^34^ or motor^35^ computations. The ANN does not employ sampling-based representations which have been proposed as possible mechanisms for probabilistic computation^36,37^, showing that explicit representation of hypotheses and uncertainty as separate regions in rate-space could serve as alternative or supplementary mechanism to sampling.

A key open question is how learning a specific environment, task, or behavioral context occurs. We observed that hypothesis coding emerges with task learning (Supplementary Fig. 6). Possible, and not mutually exclusive, mechanisms include: i) changes of the stable recurrent dynamics in RSC, as is suggested in hippocampal CA1^26^; ii) modification of dynamics by context-specific tonic inputs^1,38^; or iii) changes in how hypotheses and sensory information are encoded and read-out while maintaining attractor dynamics that generalize across environments or tasks, as indicated by the maintenance of recurrent structure across tasks in our data (Supplementary Fig. 13) and as has been shown in entorhinal^27^ and motor cortex^35^ and ANNs^39,40^, possibly helped by the high-dimensional mixed nature of RSC representations^41,42^. Further, how such processes are driven by factors such as reward expectation^43^ is an active area of research.

Our findings show that recurrent dynamics in neocortex can simultaneously represent and compute with task and environment-specific multi-modal hypotheses in a way that gives appropriate meaning to ambiguous data, possibly serving as a general mechanism for cognitive processes.

## Methods Summary

Microdrive implants with 16 tetrodes were implanted in RSC targeting layer 5, and sorted single-unit spike trains were analyzed together with mouse position and task state. All sorted neurons were included in the analysis. The ANN consisted of rate neurons with an input layer into 128 hidden recurrent units (tanh nonlinearity) into 80 output neurons, trained on random velocity trajectories in random environments of up to 4 landmarks. For the analyses in the main text, LM inputs were relayed to the ANN as a map that encoded their relative position but not identity (‘external map’ ANN, 80 input neurons). The findings were replicated with an ANN that only received binary LM presence input (‘internal map’ ANN, 11 input neurons) and non-negative ANNs (Supplementary Figs. 15-16), on a subset of environments.

## Acknowledgements

We thank Emily J. Dennis, Mehrdad Jazayeri, Kimberly Stachenfeld, and Elias Issa for comments on the manuscript. This work was supported by the NIH 1K99NS118112-01 and a Simons Center for the Social Brain at MIT postdoctoral fellowship (J.V.), National Institute of General Medical Sciences T32GM007753 (E.H.S.T), and the Center for Brains, Minds and Machines (CBMM) at MIT, funded by NSF STC award CCF-1231216 (J.P.N), and NIH R01NS106031 and R21NS103098 (M.T.H). E.H.S.T is a Paul & Daisy Soros Fellow. I.R.F. is an HHMI Faculty Scholar and this work was partially supported by awards to I.R.F. from the Office of Naval Research, the Simons Foundation through the International Brain Laboratory, and a CIFAR Senior Fellowship. M.T.H is a Klingenstein-Simons Fellow, a Vallee Foundation Scholar, and a McKnight Scholar.

## Author contributions

J.V., I.K., I.R.F. and M.T.H designed the study. I.K. and I.R.F. designed and performed the ANN component of the study. N.J.M, J.V., and J.P.N collected mouse data. J.P.N provided technical support for mouse recordings. J.V. and E.H.S.T analyzed mouse data. J.V. and M.T.H wrote the paper with input from all authors.

## Methods

### Mouse navigation behavior and retrosplenial cortex recordings

#### Drive implants

Light weight drive implants with 16 movable tetrodes were built as described before^1^. The tetrodes were arranged in an elongated array of approximately 1250×750 μm, with an average distance between electrodes was 250 μm. Tetrodes were constructed from 12.7 μm nichrome wire (Sandvik – Kanthal, QH PAC polyimide coated) with an automated tetrode twisting machine^2^ and gold-electroplated to an impedance of approximately 300 KΩ.

#### Surgery

Mice (C57BL/6 RRID: IMSR_JAX:000664) were aged 8-15 weeks at the time of surgery. Animals were housed in pairs or triples when possible and maintained on a 12-h cycle. All experiments were conducted in accordance with the National Institutes of Health guidelines and with the approval of the Committee on Animal Care at the Massachusetts Institute of Technology (MIT). All surgeries were performed under aseptic conditions under stereotaxic guidance. Mice were anesthetized with isofluorane (2% induction, 0.75–1.25% maintenance in 1 l/min oxygen) and secured in a stereotaxic apparatus. A heating pad was used to maintain body temperature, additional heating was provided until fully recovered. The scalp was shaved, wiped with hair-removal cream and cleaned with iodine solution and alcohol. After intraperitoneal (IP) injection of dexamethasone (4 mg/kg), Carprofen (5mg/kg), subcutaneous injection of slow-release Buprenorphine (0.5 mg/kg), and local application of Lidocaine, the skull was exposed. The skull was cleaned with ethanol, and a thin base of adhesive cement (C&B Metabond and Ivoclar Vivadent Tetric EvoFlow) was applied. A stainless steel screw was implanted superficially anterior of bregma to serve as electrical ground.

A 3 mm craniotomy was drilled over central midline cortex, a durotomy was performed on one side of the central sinus, and tetrode drives^1^ were implanted above Retrosplenial cortex, at around AP −1.25 to −2.5 mm and ML 0.5 mm, with the long axis of the tetrode array oriented AP, and the tetrode array tilted inwards at an angle of ∼15-20°, and fixed with dental cement. The ground connection on the drive was connected to the ground screw, and the skin around the drive implant was brought over the base layer of adhesive as much as possible to minimize the resulting open wound, sutured, and secured with surgical adhesive.

At the time of implant surgery, only two of the tetrodes were extended from the drive to serve as guides during the procedure. All other tetrodes were lowered into superficial layers of cortex within 3 days post-surgery. Mice were given 1 week to recover before the start of recordings.

#### Chronic Electrophysiology

After implant surgery, individual tetrodes were lowered over the course of multiple days until a depth corresponding to layer 5 was reached and spiking activity was evident. Data were acquired with an Open Ephys^3^ ONIX^4^ prototype system at 30kHz using the Bonsai software^5^. The tether connecting the mouse headstage to the acquisition system was routed through a commutator above the arena and was counterbalanced via a segment of flexible rubber tread.

#### Spike sorting

Voltage data from the 16 tetrodes, sampled at 30 KHz were band-pass filtered at 300-6000 Hz, and a median of the voltage across all channels that were well connected to tetrode contacts was subtracted from each channel to reduce common-mode noise such as licking artifacts.

Spike sorting was then performed per tetrode using the Mountainsort software^6^ (https://github.com/flatironinstitute/mountainsort_examples), and neurons were included for further analysis if they had a noise overlap score below 0.05, an isolation score > 0.75 (provided by Mountainsort^6^), a clear refractory period (to ensure spikes originated from single neurons), and a spike waveform with one peak and a clear asymmetry (to exclude recordings from passing axon segments), and a smooth voltage waveform and histogram (to exclude occasional spike candidates driven by electrical noise). Units were not excluded based on firing rates, tuning, or any higher order firing properties.

#### Histology

To verify the localization of the recording sites (Supplementary Fig. 3), electrolytic lesions were created by passing currents of 20 μA through a subset of tetrodes (∼4 tetrodes per animal) for 30s each under isoflurane anesthesia, and animals were perfused and brain processed 1h later. Brains were mounted with DAPI and imaged.

#### Behavioral experiment hardware

Behavior was carried out in a circular arena of 50 cm diameter. The floor of the arena was formed by a clear acrylic sheet, under which a diffusion screen and a flat-screen TV was positioned on which visual stimuli were displayed. The circular arena wall was formed by 32 flat black acrylic segments, every other one of which contained an opening for a recessed reward ports, 16 in total. Each reward port contained an optical beam break (880nm IR, invisible to mouse) that detected if a mouse was holding its nose in the port, a computer-controlled syringe pump for water reward delivery, and a dedicated beeper as a secondary reward indicator. The behavior arena was housed in a soundproof and light-insulated box with no indicators that could allow the mice to establish their heading. Video was acquired by a central overhead camera at 30 Hz using a low level of infrared light at 850 nm and the mouse position was tracked using the oat software^7^ (https://github.com/jonnew/Oat). A custom behavioral control state-machine written in Python was triggered every time a new camera frame was acquired, and the position of the animal, time passed, and port visits were used to transition the logic of the state machine (Supplementary Fig.1). For analysis purposes, all behavioral data was re-sampled to 100 Hz and synchronized to the electrophysiological data.

#### Inactivation of RSC and causal necessity for hypothesis-based computations

For pharmacological inactivation of RSC (Supplementary Fig.2), 4 mice were trained on a simplified parametric task that permitted us to causally test the role of RSC in individual recording and inactivation sessions. The task required integration of an allocentric position hypothesis with visual landmarks (Supplementary Fig. 2a,b). After mice learned the task, quantified as reaching a hit rate of above 30% in the simple conditions (high eccentricity, see Supplementary Fig.2b), they were given access to unrestricted water and implanted following the procedure described for the main experiment, but instead of a chronic drive implant, a removable cap was implanted and two burr holes were prepared above RSC and covered with dental cement (Supplementary Fig.2c). After recovery from surgery, mice were put back on water restriction over the course of one week and re-introduced to the task. Before each experiment, mice were briefly anesthetized with isoflurane, the cap was temporarily opened, and the exposed skull was wiped with lidocaine and an injection of either 50nl of 1ug/ml muscimol solution in cortex buffer per side, or the same volume of cortex solution was performed through the existing burr holes. Mice were left to recover from anesthesia for 15 min and tested on the task. Performance was assessed as the hit rate on the 1^st^ port visit per trial, and confidence level were computed via binomial bootstrap.

#### Behavioral Training

After mice had undergone preparatory surgery, they were given at least one week to recover before water scheduling began. Initially, mice received 3 ml of water per day in the form of 3 g of HydroGel (ClearH2O, Watertown, MA, USA), which was gradually reduced to 1.0–1.5 g per day. During this period, mice were handled by experimenters and habituated to the arena. Throughout the entire experiment mice were given water rewards for completion of the task and were given additional water to maintain their total water intake at 1.25-1.5ml. After initial acclimatization to the recording arena over 2 days, mice were trained on the task. Throughout the task we used white circular cues on the floor (referred to as landmarks) of ∼30mm diameter on a black background. These landmarks were the only source of light in the experiment. Mice were run every day or every other day, for a single session of 30 min to 3 hours per day. Training progressed in multiple phases:

1. Initially, mice were trained that circular visual cues on the floor of the arena indicated reward locations. One of the 16 ports was randomly selected as reward port and a cue was shown in front of this port. Visiting an incorrect port resulted in a timeout (∼1 second initially, increased later), during which the entire arena floor was switched to grey leading to a widespread visual stimulus. Visiting the correct port resulted in an audible beep from the beeper located in the port and around 0.005ml of water were delivered by the syringe pump. After a reward, a new reward port was randomly chosen, and the landmark was rotated together with the port, effectively performing a rotation of the entire task, and the next trial began. This meant that mice learned to not rely on any cues other than the visual landmark to locate the correct port. Mice usually completed this phase in by day 4.
2. We then introduced a new task phase: After each reward, the landmark disappeared and instead a blinking dot was shown in a random location in the arena. If the mouse walked over that dot, it disappeared and either a new dot in a new random location appeared, repeating the process, or the next trial was initiated. The number of required dots-chases was sampled uniformly from a range and was increased to 6-8 by the time recordings began, and the last dot was always positioned at the arena center. This task phase served to obfuscate the rotation of the task. Data acquired during this task phase was used during spike sorting but was not further analyzed. Mice learned this task phase, with 6-8 dots, by day 7 on average.
3. Throughout phases 1 and 2, we progressively introduced a requirement for the mice to hold their snouts in the reward port for increasing durations to trigger a reward or timeout. For each port visit, the required duration was randomly drawn from a uniform distribution, so on any given trial the mice did not know when exactly to expect to know the outcome of the port visit. Initially, this hold time was 500 ms, and the time range was slowly increased throughout training, depending on animal performance. By the time recordings began, a range of around 4-6 seconds was used. Mice were able to tolerate this holding time by day 20 on average.
4. Next, we introduced an identical 2^nd^ landmark at a non-rewarded port. Initially, the two landmarks were set two ports apart (e.g. ports 1 and 3), and this distance was progressively increased to 4 or 5 ports. As before, the rewarded port and landmarks were randomly rotated after each trial, but their relative positions remained stable. Visiting the reward port at the incorrect, ‘a’ landmark (and holding there for the required duration) was handled identically to visits to any other non-reward port and triggered the same time-out. As a result, mice learned to visit the ‘b’ port. Mice learned to make an initial distinction between the ports approximately by day 14-16. In one mouse we maintained this training phase until overall task performance was significant over entire sessions (Supplementary Fig.1f) but we noticed that the mouse had trouble consistently re-learning the next task phase. We therefore transitioned subsequent mice to the next phases before a stable behavior was established.
5. After the mice started learning to visit the port at the ‘b’ landmark, we introduced a view-distance limitation that made landmarks invisible from far away: The mouse’s position was tracked at 30Hz and for each landmark its brightness was modulated in real-time as a function of the mouse’s distance from it. The visibility was 0 for distances above a threshold, 1 for distances below a 2^nd^ threshold, and transitioned linearly between the two values. Initially these thresholds were set so that both landmarks were visible from the arena center (∼20 cm), then they were progressively reduced to values where at any time only one of the landmarks was visible to the mouse (∼10 cm). At this stage, mice that encounter a landmark after a new trial starts have no way of knowing whether this is the rewarded or non-rewarded landmark. Recordings began when mice were able to complete 100 trials/hour at a hit/miss rate >1. Mice reached this criterion level on average by total day 30-40 of training.

#### Statistical Analysis

Analyses were carried out using custom code in Matlab (Mathworks). Unless stated otherwise, confidence intervals were computed at a 95% level via bootstrap, and p-values were computed using a Mann–Whitney U test or Wilcoxon signed-rank test. In figures, significance values are indicated as ‘NS’ (P>0.05), ‘*’ (P ≤ 0.05), ‘**’ (P ≤ 0.01) or ‘***’ (P ≤ 0.001).

#### Behavior Analysis

Recording sessions were included once mice performed the task well enough to achieve a session average hit/miss ratio > 1, indicating that mice could infer the correct port between the ‘a’ and ‘b’ landmarks (a correct rate of >1/16 would indicate that they can associate landmarks with rewarded ports, but not that they can infer landmark identity). Because landmarks are only visible sequentially after full training, a ratio >1 shows that mice employed a memory based strategy where they used a prior hypothesis derived from seeing or not seeing the 1^st^ landmark to infer the identity of the 2^nd^ landmark they encounter. Only sessions with at least 50 recorded single neurons, and with at least 50 minutes of task performance were included. This yielded 16 sessions from 4 mice. For some analyses, particularly for analyses where trajectories of the mice were matched across trial types to control for potential motor and sensory confounds, additional selection criteria were applied yielding a lower N of sessions that could be used, this is stated for the respective analyses. For plots of the learning rates we included trials where mice encountered their 1^st^ landmark after 20 seconds or faster to exclude periods where mice were not engaged, plots using all trials are also included in the supplement.

#### Behavioral epochs

For analysis, each trial was split into epochs: The time between the onset of a trial (right after the mouse completes the preceding re-initialization procedure) and the onset of the reward (the first time the mouse could know whether it reached the correct port, other than by process of elimination after visiting other ports) was split up based on the amount of information the mouse could have accumulated: the initial state when mice had not seen any landmark was labeled ‘LM0’, time after the first landmark encounter was labelled ‘LM1’, and after the 2^nd^ as ‘LM2’. The timepoints when landmarks became visible and the mouse transitioned from LM0 to LM1 or from LM1 to LM2, referred to as ‘landmark encounters’ were defined as the timepoint when landmark visibility exceeded 50%.

For analyses of the correlation of neural state and eventual behavioral outcomes, each 2^nd^ landmark encounter was further categorized as whether it occurred at the ‘a’ or ‘b’ landmark. For behavioral analyses in Fig.4d, trials were further categorized by whether they led to a correct port visit or to a incorrect visit and a time-out.

#### Similarity of spatial tuning across conditions

Changes in spatial tuning in individual RSC neurons as mice encounter successive landmarks (Fig.1f) was quantified by the Euclidian distance of their spatial tuning profiles (in an 8×8 map, for each comparison non-visited ties were omitted). As an internal control, distance between tuning profiles within –condition and across-condition were compared using non-overlapping 1 minute segments. For each comparison (LM1 vs. LM2 and LM0 vs. LM1), the split spatial tuning maps were compared either within the conditions, e.g. within LM1 and within LM2, and compared to distances between LM1 and LM2 maps.

#### Neural decoding of mouse position

To decode the mouse position from RSC firing rates, neural firing rates were first low-pass filtered at 1Hz with a single-pole butterworth filter. The resulting firing rate time series were used to predict the mouse position as 100 categorical variables forming a 10×10 bin grid, (bin width = 50 mm). The network was made up of a single LSTM layer with 20 units, and a fully connected layer into a softmax output into the 100 possible output categories. For analyses of intermediate information content of the decoder, the network input into the final softmax layer was analyzed.

Decoding was re-initialized for each trial. For each decoded trial, all other trials served as training set. For analysis of how the neural coding of position was dependent on the LM state of the mouse (Supplementary Fig. 5a), the same analysis was repeated with training and testing data further divided by LM state. For analysis of the decoding performance, the output likelihood from the decoder was evaluated at the mouse’s true position for all positions that were shared across conditions for this session. Statistical analysis was then performed on a per-session average likelihood (not weighted by number of trials per session).

#### Neural decoding of LM state

For the analysis of LM state (Fig.1e), trials with at least 0.5 seconds of data from all 3 states were used (16 sessions, 486 total trials), and individual trials were held out from training for decoding. Firing rates were low-pass filtered with a causal single-pole butterworth filter at 0.05 Hz, and LM state (0, 1 or 2) was decoded independently for each timepoint using a categorical linear decoder (dummy variable coding, (N_neurons_ +1)*3 parameters), or a neural network with no recurrence, using a single 20 unit layer receiving instantaneous firing rates, into a 6 unit layer, into 3 softmax outputs.

#### Dimensionality analysis

(Supplementary Fig. 13c). Principal component analysis was performed by first computing the covariance matrices of the low-pass filtered (as before) firing rates, and plotting their eigenvalue spectra, normalized by sum. Each scaled eigenvalue corresponds to a proportion of explained variance. Spectra are plotted together with a control spectrum computed from covariances of randomly shuffled data. For a description of the method used to compute the correlation dimension of RSC rates (Supplementary Fig. 13d) see the heading ‘Correlation dimension’ in the section about artificial neural network methods below.

#### Prediction of firing rates across RSC population

For quantification of the independence of individual RSC neurons from the surrounding RSC population (Supplementary Fig. 13f,g), the firing rates of each neuron were predicted from those of all other neurons using linear regression. Rates were first filtered at 0.01-0.5 Hz with a 3^rd^ order butterworth filter, and sub-sampled to 3.3 Hz. Each neuron’s rate was predicted with L_1_ regularized linear regression ^8^ (lasso, *λ* ≈ 0.0001) from the rates of all other neurons and preceding firing rates using 8 lags (∼0.2.5 sec). Goodness of fit was quantified as the proportion of variance explained: 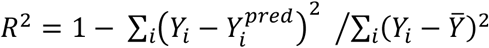. Predictions were computed both within condition (LM1, LM2, and dot-hunting phase), as well as across conditions, where the model was fit using coefficients determined from the other conditions.

#### Computation of firing rate distribution entropies

Entropies of empirical firing rate distributions were computed in bits via their Shannon entropy: 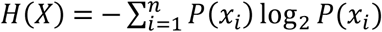, relative to a uniform histogram of the same size: *Ĥ*(*X*) = −(*H*(*X*) − *H*(*uniform*)). In cases where zeros appeared, a small offset term <<1 was added and all histograms were normalized to a sum of 1. For example, *Ĥ*([1 0]) = *Ĥ*([1 1 0 0]) = 1 *bit* and *Ĥ*([1 1 1 1.3]) ≅ 0.01 *bit*.

#### Trial-to-trial variance of firing rates conditioned on position

For analysis of whether partial hypothesis representation in the LM1 state corresponds to trial-by-trial changes in firing rates, evident in bimodal firing rate histograms, histograms of hidden unit firing rates of the ANN, conditioned on binned 1-D position are displayed (Supplementary Fig. 10a). Data are from Experiment configuration 2 (See Methods, section ‘Overview over experiment configurations used with ANNs’). Tuning curves were calculated using 20 bins of location/displacements and normalized individually for each neuron. The first time step in each trial and time steps with non-zero landmark input were excluded from the analysis. For histograms, each condition was binned in 100 column bins and neuron rates in 10 row bins. Histogram were normalized to equal sum per column. For analysis of RSC firing rates (Supplementary Fig. 10b-d), we did not observe bimodal rate distributions and instead quantified the dispersion of the rate distributions via their entropy: Firing rates were low-pass filtered at 0.5 Hz to bring them into the time scale of navigation behavior, and firing rate histograms were computed with 8 bins spanning from each neurons lowest to highest firing rate per neuron, for each spatial bin in a 4×4 grid. Because the computation of histogram entropy is biased by the N of samples, for each spatial bin, the same number of time points were used for the LM1 and LM2 conditions. The dispersion of the firing rate distribution was then computed as average entropies per cell across all space bin, and compared across the two conditions.

#### Analysis of encoding of angular position and displacement from last seem landmark

Firing rate profiles were analyzed in two reference frames: Global angle of the mouse in the arena, and relative angle to the last visible landmark. Only time-points from the foraging state where the distance of mouse from the center of the arena exceeded 70% of the arena diameter were included. Time points from the LM1 and LM2 conditions were sub-sampled to yield matched N of time points. Firing rates were analyzed in a –π to π range in 6 bins by computing their entropy as described before.

#### Pairwise correlation of firing rates

Recordings were split into LM[0,1,2] states as before, firing rates were low-pass filtered at 1Hz, and the Pearson correlation coefficient between each pair of neurons was computed. For display purposes, the neurons were re-ordered by first computing the matrix for the LM1 state, applying hierarchical clustering^9^, and the resulting re-ordering was applied to both LM1 and LM2 conditions. This re-ordering has no impact on any further analyses. For summary statistics, we computed the correlation of correlations for each session.

#### Low-dimensional embedding of neural activity

Neural firing rates were band-pass filtered as before, and an initial smoothing and dimensionality-reduction step was performed by training a small LSTM with a single layer of 30 units to decode the mouse position. The hidden unit activations were then embedded in 3-D space with the isomap algorithm^10^, using the Toolbox for Dimensionality Reduction by Laurens van der Maaten^11^.

#### Analysis of speed of neural state evolution

(Supplementary Fig. 14a) For quantification of how fast the neural state evolves, the firing rates of the entire population were computed by low-pass filtering the spike trains at 1Hz (3^rd^ order butterworth filter), and the speed of the 5 largest principal components of the resulting vector in Hz/sec were related to the running speed of the mouse (m/sec, also low-pass filtered at 1Hz), or the change in landmark brightness (percent/sec). Data were binned in 30 bins from 0-0.5m/sec and 10 bins from 0.5-2 m/sec for running speed, and 10 bins from −50 to 50% and 10 bins for ±50-200%. Confidence intervals were computed by treating median data from each session as independent samples.

#### Analysis of context-encoding in RSC across similar motor and sensory states

To study the encoding of context with minimal sensory and motor confounds (Fig. 4, and Supplementary Fig. 14), we split the appearances of the 2^nd^ landmark into two groups depending on whether the 2^nd^ landmark is ‘a’ or ‘b’, as described in the main text. We then manually selected subsets of trials where egocentric paths just before the appearance of the 2^nd^ landmark are matched across the two groups. Fig. 4a shows an example of such matched approach paths/trials. Sessions in which at least 16 trials could be matched were used for these analyses, yielding a total of 133 trials from 6 sessions (per session: 16, 23, 24, 24, 25 and 21). For each session, all of these trials were aligned to the time when the 2^nd^ landmark became visible, yielding a set of time ranges where the animals experienced similar visual inputs, performed similar locomotion behavior, but potentially encoded different prior experience leading them to subsequently disambiguate the perceptually identical 2^nd^ landmark as ‘a’ or ‘b’.

To test whether there was consistent encoding of this context in RSC, we then compared the distances across these groups in 3-dimensional neural activity space (see ‘Low-dimensional embedding of neural activity’) to distances within the groups (Fig.4d, Supplementary Fig.14). This test was performed at the point where the 2^nd^ landmark became visible to assess encoding of prior context, as well as 200ms afterwards to assess how the identity of the (now visible) landmark affects encoding in RSC.

#### Analysis of smooth neural trajectories across sessions

(Fig.4c, Supplementary Fig.14) To assess if neural trajectories were determined by population dynamics that were stable across trials and could therefore serve as substrate for the computation performed by the mice, we tested whether neural trajectories behaved consistent with a laminar flow regime where neighboring particles (in our case neural firing rate vectors) remain neighbors for a significant amount of time, or whether they decorrelate quickly (Fig.4c, Supplementary Fig.14e,f). To assess temporal dynamics of the neural spiking without imposing any smoothing, we investigated raw spike counts in 750ms windows for this analysis. For each session, an initial set of pairwise high-dimensional distances in spike-counts between the trials with egocentrically similar paths (see prev. section) was computed from the last 750ms preceding the appearance of the 2^nd^ landmark. These distances were then correlated with those in a second sliding window, Supplementary Fig. 14f). An offset of 0 seconds was defined as the point where both windows stopped overlapping. The correlation coefficient R was then computed for increasing window offset up to 2 seconds. Summary statistics were computed across sessions by first shifting each session individually by its 95% level for R (from a shuffled control) which results in the summary plot showing a highest value for R of ∼0.8 even for offsets where the windows fully overlap and the uncorrected R value is 1. Because of this offset, the null level for each trial is now at R=0. We then computed the CIs for the group via bootstrap relative to this level.

#### Analysis of direction of neural trajectories

(Supplementary Fig. 14g,h) To further test if neural trajectories were determined by population dynamics that were stable across trials, and were independent of the interpretation of the 2^nd^ (locally ambiguous) landmark, we tested if neural activity evolved in similar directions across trials if it started close together in 3-dimensional neural activity space (see ‘Low-dimensional embedding of neural activity’). We therefore looked at neural trajectories within the motor and sensory-matched LM2 approaches where the neural state at the point where the 2^nd^ landmark became visible started neurally close to other trials from the opposing class. For example, for a LM2_a_ trial, we looked if this trial might follow other close-by LM2_b_ trials. We computed neural proximity in the 3-dimensional neural embedding (see above) and defined close-by trials as ones that were within 1 arbitrary units in Euclidean distance in the isomap embedding around the time when the 2^nd^ landmark became visible, yielding a total of 42 out of 79 trials with close neighbors from opposing classes from the 5 sessions, one session was excluded because the neural activity in the relevant time ranges was collapsed onto a point in the LSTM embedding. As a control, we also selected corresponding neurally furthest points. Similarity of neural evolution was then quantified as the angular difference between the trials in (3-dimensional) LSTM space over time, in order to assess co-evolution independently of the initial selection by distance. Significance was computed via bootstrap across trials, vs random alignments corresponding to a 90 degree difference.

#### Behavior prediction

For the behavior prediction analysis, sessions with at least 5 correct and incorrect port visits after the 2^nd^ port visit were used (N=11) and an equal number of hit and miss trials (outcome of next port visit is a timeout or a correct) were selected, leading to a chance prediction level of 0.5. The spike rates from the 5 seconds preceding the 2^nd^ landmark becoming visible, binned into one-second bins, were used to predict the behavioral outcome with a binary classification decision tree with a minimum leaf size of 6, previously determined via cross-validation. Predictions for each trial were fit using all other trials.

#### Specificity of landmark encounter coding to the foraging task

(Supplementary Fig. 6) We trained a decoder to predict either the number of encountered dots in the main task, or in the dot-hunting task. These tasks were interleaved and the same neurons were used. Train and test sets were split by trial, and decoding was performed with a regression tree on lowpass-filtered firing rates as before, performance was quantified as mean error on the N of landmarks. Only the first 2 landmarks were predicted in the dot-hunting task to allow use of the same classifier across both. Decoding performance was compared between the within-class (e.g. decode main task encounters with decoder trained on other trials in the main task) and cross-class (e.g. decode dot-hunting from decoder trained on the main task etc.).

#### Analysis of neural coding as a function of task performance

(Supplementary Fig. 6) To test whether the encoding of hypothesis states in RSC is specific to task performance, we analyzed a larger number of sessions from the entire period during which two landmarks with local visibility were used (92 recording sessions in total). We analyzed the effect of task performance on the behavior prediction analysis (as described above, and in Supplementary Fig. 6). We also analyzed the more general decoding of landmark encounter count (same method as’ Specificity of landmark encounter coding to the foraging task’ or Fig.1) in all of the 92 sessions with 2 landmarks, and correlated decoding performance with task-performance on a per-session level. As an additional control, we performed the same analysis on the N of dots encountered in the interleaved dot-hunting task. For all of these analyses analogous method as for the non behavior-correlated analyses was used.

#### Artificial neural networks

A simulated animal runs with varying velocity in a circular environment starting from a random, unknown position and eventually infers its position using noisy velocity information and two, three or four indistinguishable landmarks. A trial consists of a fixed duration of exploration in a fixed environment, starting from an unknown starting location; the environment can change between trials. Environments are generated by randomly drawing a constellation of 2-4 Landmarks, and the network must generalizably localize in any of these environments when supplied with its map. The network must adjust its spatial inference computations on the basis of the configurations of the different environments, without changing its weights; thus, the adjustments must be dynamic. In the internal map scheme (Supplementary Fig 15), an input cell simply encodes by its activation whether the animal is at any landmark; it does not specify the location of the landmark, the identity of the environment, or the spatial configuration of the various landmarks in the environment. The task in the internal map scheme is substantially harder, since the network must infer the configuration of landmarks in the environment purely from the time sequence of landmark visits, while simultaneously localizing itself within the environment. Information about the maps must be acquired and stored within the network. To make the task tractable, we limit training and testing in the internal map setting to four specific environments.

In the external map task (Figs. 2,3, Supplementary Figs. 6-12), landmark locations were random and the set of locations (map) were provided to the network, whereas in the internal map task (Supplementary Fig. 15) one of four landmark configurations was used, but the maps were not provided to the network. landmarks could only be observed a short distance. A three-layer network with a recurrent hidden layer was trained to infer location. Velocity and landmark encounter information were encoded in the input layer, and all weights of the network were trained. The training target for the output layer was activation of a unit with von Mises tuning and preferred location matching the true location.

Network performance was compared to a number of alternative algorithms: Path integration + correction integrated the noisy velocity information starting from an initial location guess and corrected this estimate by a reset to the coordinates of the nearest landmark when a landmark was encountered. Particle filters approximated sequential Bayesian inference given the available velocity and landmark information, with each particle capturing a location hypothesis whose posterior probability is given by an associated weight. Particle locations are updated using velocity information and particles are reweighted after landmark encounters. The enhanced particle filter also reweights particles when a landmark is expected but not encountered, thus can infer location not only from the presence but also from the absence of landmarks. The output and hidden representations of the trained network were evaluated in a variety of conditions involving both random and fixed landmark locations and trajectories with random and fixed velocities.

#### Definition of environments and trajectories

The task is defined by a simulated animal moving along a circular track of radius 0.5 m for 10 seconds. The animal starts at a random, unknown position along the circle at rest and starts running along a trajectory at non-constant velocity. A trajectory is sampled every *dt* = 0.1s in the following way: At each time t, acceleration at is sampled from a zero-mean Gaussian distribution with standard deviation σ_a_ = π/4 m/s^2^ that is truncated if |a_t_| > π/2 *m/s*^2^. Acceleration is integrated to obtain the velocity v_t_ and truncated if |v_t_| > v_max_ = π/2 *m/s*. The actual location on the track is the integral of this velocity.

In a trial of the external map task, the locations of *K* = 2, 3, or 4 indistinguishable landmarks were determined sequentially: the first landmark was sampled from a uniform random distribution on the circle, with subsequent landmarks also sampled from a uniform random distribution but subject to the condition that the minimum angular distance from any previously sampled landmark is at least δ = π/9 rad.

The internal map task involved four environments, each with a unique configuration of landmarks: two environments had two landmarks, one had three and the last had four. Landmark locations in the four environments were chosen so that pairwise angular distances were sufficiently unique to allow the inference of environment identity. Landmark coordinates in environment ei were given by: *e*_1_ = {0, 2π/3} rad, *e*_2_ = {1.9562, 3.7471} rad, *e*_3_ = {0.2641, 1.2920, 3.7243} rad, *e*_4_ = {3.0511, 3.8347, 5.1625, 5.7165} rad.

#### Experiment configurations used with ANNs

After training, the networks were evaluated in different testing configurations that each consisted of a distribution over landmark configurations and trajectories:

*Experiment configuration 1*. Training distribution: This test set was generated exactly as in the training set, as described in section “Definition of environments and trajectories”. 5 different

*Experiment configuration 2*. Fixed landmarks, random trajectories: The landmark configuration was given by two landmarks located at *e* = {0, 2π/3}, the trajectories were sampled in an identical way as in the training distribution. Note that this landmark configuration corresponds to the first environment in the internal map task.

*Experiment configuration 3*. Fixed landmarks, constant velocity trajectories: The landmark configuration was given by two landmarks located at *e* = {0, 2π/3} and the trajectories were given by constant velocity trajectories with |v_t_| = v_max_/2. The initial position and the direction of the trajectory was random.

*Experiment configuration 4*. Two variable landmarks, constant velocity trajectory: The landmark configuration was given by two landmarks located at *e* = {0, 2π/3 + απ/3}, where α ∈ [0, 1]. The trajectories were given by constant velocity trajectories with |v_t_| = v_max_/2 and the initial position and the direction of the trajectory was random.

*Experiment configuration 5*. Two environments, random trajectories: The landmark configuration was given by either *e*_1_ or *e*_2_ of the internal map task, trajectories are random.

#### Landmark observation

The animal is considered to have encountered a landmark if it approached within *d*_*min*_ = *v*_*max*_ · *dt/2* = π/40 *m*/2 = π/20 rad. This threshold is large enough to prevent an animal from “missing” a landmark even if it is running at maximum velocity. This ‘visibility radius’ is smaller than the one we used for the mouse behavior experiments (Fig.1). In the ANN experiments, landmark encounters were therefore roughly coincident with the agent’s position coinciding with the landmark, whereas in the mouse data, landmark encounters occur a significant distance away from the landmark, when it becomes visible (e.g. Fig.4a). In the same way as in the mouse behavior analysis, hovering around the same landmark or approaching the same landmark consecutively would only trigger a landmark encounter at the first approach; a new encounter was only triggered if the animal approached a landmark different than the previous one, equivalent to the definition used in the analysis of mouse behavior. Also, only trials in which the animal encountered at least two different landmark were included.

#### Sensory noise

The largest sources of uncertainty in the tasks were the unknown starting position and the indistinguishability of the landmarks. In addition, we assumed that the velocity information and the landmark-location memory (in the external map scenario) were corrupted by noise. At each time step of size *dt* = 0.1, the velocity input to the network corresponded to the true displacement *vdt* corrupted by zero-mean Gaussian noise of standard deviation σ_v_ = *v*_*max*_*dt*/10. In the external map task, the landmark map provided to the network and particle filter was corrupted by zero-mean Gaussian noise with standard deviation *σ*_*l*_ = π/50 rad, without changing the relative landmark positions: The map was coherently slightly rotated at a landmark encounter, and the rotation was independently sampled at each landmark encounter.

#### ANN preferred firing at landmark locations

(Supplementary Fig. 5c) This analysis was performed by evaluating the network of the external map task on the experiment configuration 1 of the internal map task. First, location tuning curves were determined after the second landmark encounter using 5000 trials from distribution 1 and using 50 location bins. Tuning curves were calculated separately for each of the four environment of the internal map task. Preferred location was determined to be the location corresponding to the tuning curve maximum. The density of preferred locations smaller than distance *d*_min_ away from a landmark was then compared to the density of preferred locations further away from landmarks.

#### Network architecture and training

The network consisted of three layers of rate neurons with input-to-hidden, hidden-to-hidden and hidden-to-output weights. All weights were trained.

##### Network input

The input layer consisted of 80 neurons in the external map case and 11 neurons in the internal map case. Ten neurons coded for velocity corrupted by noise (noise as described above). The velocity neurons had a minimum firing rate between 0 and .2 and a maximum firing rate between .8 and 1 in arbitrary units, and within this output range coded linearly for the whole range of velocity between −*v*_*max*_ and *v*_*max*_. Negative and positive velocity here corresponds to clockwise and counterclockwise travel respectively.

The remaining neurons (70 in the external map case and 1 in the internal map case) coded for landmark input and were activated only at the time step of, and up to three time steps after an landmark encounter. In the external map case, the landmark input simultaneously encoded the locations of all landmarks in the environment, thus supplying a map of the environment, but contained no information about which landmark was currently encountered. The landmark neurons had von Mises tuning with preferred locations *x*_*j*_ = (*j* − 1) · 2π/70 rad, *j* = 1…70, that tiled the circle equally. Given *n* landmarks at locations *l*_*i*_, i = 1…*n*, the firing rate of the *j*-th landmark input neuron was given by

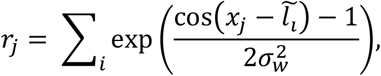

where 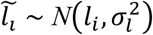 is the noise-corrupted landmark coordinate (see “Sensory noise” section). This mixture of von Mises activation hills produces the pattern depicted as the “map” input in Supplementary Fig.7a.

In the internal map case (Supplementary Fig. 15), the landmark input neuron consisted of a single binary neuron that responded for four time steps with activation 1 in arbitrary units whenever a landmark was encountered. This input encoded neither environment identity nor landmark location.

##### Hidden layer

The hidden layer consisted of 128 recurrently connected neurons. The activation **h**_*t*_ of hidden layer neurons at time step t was determined by **h**_*t*_ = tanh(*W*_*x*_**x**_*t*_ + W_*h*_**h**_*t*_−1 + **b**), where **x**_*t*_ are the activations of input neurons at time step t, *W*_*x*_ are the input-to-hidden weights, *W*_*h*_ are the hidden-to-hidden weights and **b** are the biases of hidden neurons. The nonlinearity should be considered as an effective nonlinearity at long times; since the time step *dt* = 0.1s was large compared to a typical membrane time constant (τ ≈ 0.02s), we did not include an explicit leak term.

##### Hidden layer (non-negative network)

In the non-negative network (Supplementary Fig. 16), the recurrent activation was determined by ***h***_*t*_ = tanh([*W*_*x*_**x**_*t*_ + W_*h*_**h**_*t*−1_ + **b**]_+_), where [*u*]_+_ denotes rectification.

##### Output layer

The output layer consisted of a population of 70 neurons with activity **o**_*t*_ given by **o**_*t*_ = tanh(*W*_*o*_**h**_*t*_ + **b**_*o*_), where *W*_*o*_ are the output weights and **b**_*o*_ the biases of the output neurons.

##### Network training

The training targets of the output layer were place cells with von Mises tuning of width σ_*o*_ = π/6 rad to the true location *y*_*t*_,

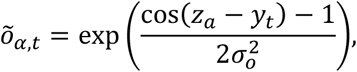

where *z*_*α*_, *α* = 1…70 are the equally spaced preferred locations of each training target.

The network was trained by stochastic gradient descent using the Adam algorithm^12^, to minimize the average square error between output ***o***_*t*_ and training targets 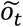, with the average taken over neurons, time within each trial, and trials. The gradients were clipped to 100. The training set consisted of 10^6^ independently generated trials. During training, performance was monitored on a validation set of 1000 independent trials and network parameters with the smallest validation error were selected. All results were cross-validated on a separate set of test trials to ensure that the network generalized across new random trajectories and/or landmark configurations.

##### Network location estimate

Given the activity of the output layer at time t, we define the network location estimate for that time to equal the preferred location (the preferred location was set over training) of the most active output neuron:

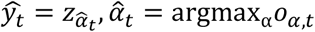

#### Performance comparisons

In Figure 2b, we compared the performance of the network in the external map task with a number of alternative algorithms. To ensure a fair comparison, we make sure that each alternative algorithm has access to exactly the same information as the network: the landmark identities are indistinguishable and both velocity and landmark location information are corrupted by the same small amount of sensory noise. Error statistics are computed from 5000 trials.

##### Path integration + correction

This algorithm implements path integration and landmark correction using a single location estimate, similar to what is implemented in hand-designed continuous attractor networks that implement resets at boundaries or other landmarks [37, 36, 25, 9]. The algorithm starts with an initial location estimate at y = 0 (despite the true initial location being random and unknown), and integrates the noise-corrupted velocity signal to obtain location. At each landmark encounter the algorithm corrects its location estimate to equal the coordinates of the landmark nearest to its current estimate.

##### Basic Particle filter

Particle filters implement approximate sequential Bayesian inference using a sampling-based representation of the posterior distribution. Here, the posterior distribution over location at each time point is represented using a cloud of weighted particles, each of which encodes through its weights a belief, or estimated probability, of being at a certain location. In the beginning of the trial, *N*_*p*_ = 1000 particles are sampled from a uniform distribution along the circle and weighted equally. In the prediction step, particles are independently propagated using a random walk whose mean is the noise-corrupted velocity update and whose standard deviation is the velocity noise *σ*_*v*_. In the absence of a landmark encounter, particle weights remain unchanged and the particle cloud diffuses. If a landmark is encountered, the importance weights *w*_*t,β*_ of particles *β* = 1…Np are multiplied by

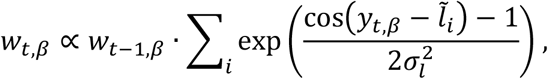

where *y*_*t,β*_ are the current estimates of the particles, and the weights are subsequently normalized such that 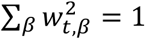. If the effective number of particles becomes too small, i.e. 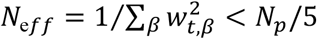, the particles are resampled using low variance sampling^13^ and the weights equalized. This resampling step both allows for better coverage of probabilities and permits the particle cloud to sharpen again. The particle filter estimate at a given time point is given by the weighted circular mean *ŷ*_*t*_ = arg(∑_*β*_ *w*_*t,β*_exp(*iy*_*t,β*_)) of the particle locations. In addition we also calculate the circular variance as var(*y*_*t*_) = 1 − |∑_*β*_ *w*_*t,β*_ exp(*iy*_*t,β*_)|.

##### Enhanced particle filter

This particle filter has identical initialization, prediction step and weight update at landmark encounters as the basic particle filter and proceeds in exactly the same way until the first landmark encounter. Subsequently, the enhanced particle filter can also use the absence of expected landmark encounters to narrow down its location posterior, similar to the network’s ability shown in Supplementary Fig. 7. This is implemented in the follow way: If a particle comes within the observation threshold *δ* of a possible landmark location but no landmark encounter occurs, the particle is deleted by setting its weight to zero; afterwards the particle weights are renormalized. A complication to this implementation is that a subsequent landmark encounter only occurs if the current landmark is different than the previous one (see section “Landmark encounters”); to prevent the deletion of particles that correctly report a landmark at the current position but do not receive a landmark encounter signal because it is the same landmark as previously encountered, particles are only deleted if they come within the observation threshold *δ* to a possible landmark that is different than the last landmark and do not encounter it. In case all particles have been deleted, particles are resampled from a uniform distribution and their weights are equalized. As for the basic particle filter, particles are resampled whenever the effective number of particles becomes too small 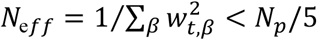. Also the particle filter estimate ŷ_*t*_ = arg(∑_*β*_ *w*_*t,β*_exp(*iy*_*t,β*_)) and the circular variance var(*y*_*t*_) = 1 − |∑_*β*_ *w*_*t,β*_ exp(*iy*_*t,β*_)| is calculated in an identical way.

#### Analysis of location disambiguation in output layer

The timing and accuracy of location disambiguation in Supplementary Fig. 7 was calculated in the following way: We first constructed the trajectory of the “alternative location hypothesis”, corresponding to the location estimates of a model animal that made the wrong location disambiguation at the first landmark encounter, but otherwise updated its location by the correct velocity. This trajectory is shifted relative to the true trajectory by a constant distance equal to the distance between the two landmarks. At each point in time, we then identified the two neurons in the output population whose preferred locations were closest to that of the true and alternative trajectory, respectively; the activation of these neurons roughly corresponded to the height of the activation bump corresponding to the true and alternative location hypothesis as seen in Supplementary Fig. 7c&d. The disambiguation time was defined as the earliest time after which either the true or alternative location bump height fell below a threshold of 0.1 and stayed beyond that threshold until the end of the trial. To determine the accuracy of location disambiguation the network estimate at the last landmark interaction was analyzed. If this network estimate was closer to the true than to the wrong landmark location the trial was categorized as a correct trial, otherwise it was categorized as an incorrect trial.

#### State space analysis

We performed principal components analysis (PCA) on the hidden neuron states from training trials to obtain the top three principal directions. We then projected network states obtained from the distribution of testing trials 2 or 3 (see SI) onto these principal directions. The resulting reduced-dimension versions of the hidden neuron states from testing trials are shown in Fig.2 and Supplementary Figs. 8, 12, and 15.

#### Correlation dimension

To calculate the correlation dimension for the ANN and RSC activity we first performed linear dimensionality reduction (PCA) on hidden layer activations from the training trials, retaining 20 principal components. For RSC data, rates were low-pass filtered at 0.5Hz first. In the 20-dimensional space, we randomly picked 1000 base points (500 for RSC). From each of these base points, we estimated how the number of neighbors in a ball of radius R scales with R. The minimum ball radius was determined such that the logarithm of the number of neighbors averaged over base points was near 1. The maximum radius was set to 10 times the minimum radius, and intermediate values for the radius were equally spaced on a log-scale. The slope of the linear part of the relationship between the logarithm of number of neighbors versus ball radius determined the fractal dimension

## References

1. Mante, V., Sussillo, D., Shenoy, K. V. & Newsome, W. T. Context-dependent computation by recurrent dynamics in prefrontal cortex. Nature 503, 78–84 (2013).

2. Sarafyazd, M. & Jazayeri, M. Hierarchical reasoning by neural circuits in the frontal cortex. Science 364, (2019).

3. Vyas, S., Golub, M. D., Sussillo, D. & Shenoy, K. V. Computation Through Neural Population Dynamics. Annu. Rev. Neurosci. 43, 249–275 (2020).

4. Smith, R. C. & Cheeseman, P. On the Representation and Estimation of Spatial Uncertainty. Int. J. Robot. Res. 5, 56–68 (1986).

5. Cho, J. & Sharp, P. E. Head direction, place, and movement correlates for cells in the rat retrosplenial cortex. Behav. Neurosci. 115, 3–25 (2001).

6. Alexander, A. S. & Nitz, D. A. Retrosplenial cortex maps the conjunction of internal and external spaces. Nat. Neurosci. 18, 1143–1151 (2015).

7. Voigts, J. & Harnett, M. T. Somatic and Dendritic Encoding of Spatial Variables in Retrosplenial Cortex Differs during 2D Navigation. Neuron 105, 237–245.e4 (2020).

8. Mao, D., Kandler, S., McNaughton, B. L. & Bonin, V. Sparse orthogonal population representation of spatial context in the retrosplenial cortex. Nat. Commun. 8, (2017).

9. Murakami, T., Yoshida, T., Matsui, T. & Ohki, K. Wide-field Ca2+ imaging reveals visually evoked activity in the retrosplenial area. Front. Mol. Neurosci. 8, (2015).

10. Fischer, L. F., Mojica Soto-Albors, R., Buck, F. & Harnett, M. T. Representation of visual landmarks in retrosplenial cortex. eLife 9, e51458 (2020).

11. Voigts, J., Newman, J. P., Wilson, M. A. & Harnett, M. T. An easy-to-assemble, robust, and lightweight drive implant for chronic tetrode recordings in freely moving animals. J. Neural Eng. (2020) doi:10.1088/1741-2552/ab77f9.

12. Burak, Y. & Fiete, I. R. Accurate Path Integration in Continuous Attractor Network Models of Grid Cells. PLOS Comput. Biol. 5, e1000291 (2009).

13. Samsonovich, A. & McNaughton, B. L. Path Integration and Cognitive Mapping in a Continuous Attractor Neural Network Model. J. Neurosci. 17, 5900–5920 (1997).

14. Widloski, J. & Fiete, I. R. A Model of Grid Cell Development through Spatial Exploration and Spike Time-Dependent Plasticity. Neuron 83, 481–495 (2014).

15. Hardcastle, K., Ganguli, S. & Giocomo, L. M. Environmental Boundaries as an Error Correction Mechanism for Grid Cells. Neuron 86, 827–839 (2015).

16. Hollup, S. A., Molden, S., Donnett, J. G., Moser, M.-B. & Moser, E. I. Accumulation of Hippocampal Place Fields at the Goal Location in an Annular Watermaze Task. J. Neurosci. 21, 1635–1644 (2001).

17. Lee, I., Griffin, A. L., Zilli, E. A., Eichenbaum, H. & Hasselmo, M. E. Gradual Translocation of Spatial Correlates of Neuronal Firing in the Hippocampus toward Prospective Reward Locations. Neuron 51, 639–650 (2006).

18. Nieh, E. H. et al. Geometry of abstract learned knowledge in the hippocampus. Nature 1–5 (2021) doi:10.1038/s41586-021-03652-7.

19. Sleezer, B. J., Castagno, M. D. & Hayden, B. Y. Rule Encoding in Orbitofrontal Cortex and Striatum Guides Selection. J. Neurosci. 36, 11223–11237 (2016).

20. Panichello, M. F. & Buschman, T. J. Shared mechanisms underlie the control of working memory and attention. Nature 1–5 (2021) doi:10.1038/s41586-021-03390-w.

21. Scott, B. B. et al. Fronto-parietal Cortical Circuits Encode Accumulated Evidence with a Diversity of Timescales. Neuron 95, 385–398.e5 (2017).

22. Harvey, C. D., Coen, P. & Tank, D. W. Choice-specific sequences in parietal cortex during a virtual-navigation decision task. Nature 484, 62–68 (2012).

23. Yang, T. & Shadlen, M. N. Probabilistic reasoning by neurons. Nature 447, 1075–1080 (2007).

24. Odoemene, O., Nguyen, H. & Churchland, A. K. Visual evidence accumulation behavior in unrestrained mice. bioRxiv 195792 (2017) doi:10.1101/195792.

25. Xue, C., Kramer, L. E. & Cohen, M. R. Dynamic task-belief is an integral part of decision-making. bioRxiv 2021.04.05.438491 (2021) doi:10.1101/2021.04.05.438491.

26. Guo, W., Zhang, J. J., Newman, J. & Wilson, M. A. Latent learning drives sleep-dependent plasticity in distinct CA1 subpopulations. bioRxiv 2020.02.27.967794 (2020) doi:10.1101/2020.02.27.967794.

27. Yoon, K. et al. Specific evidence of low-dimensional continuous attractor dynamics in grid cells. Nat. Neurosci. 16, 1077–1084 (2013).

28. Gardner, R. J. et al. Toroidal topology of population activity in grid cells. bioRxiv 2021.02.25.432776 (2021) doi:10.1101/2021.02.25.432776.

29. McKenzie, S. et al. Preexisting hippocampal network dynamics constrain optogenetically induced place fields. Neuron 109, 1040–1054.e7 (2021).

30. Banerjee, A. et al. Value-guided remapping of sensory cortex by lateral orbitofrontal cortex. Nature 585, 245–250 (2020).

31. Inagaki, H. K., Fontolan, L., Romani, S. & Svoboda, K. Discrete attractor dynamics underlying selective persistent activity in frontal cortex. bioRxiv 203448 (2017) doi:10.1101/203448.

32. Finkelstein, A. et al. Attractor dynamics gate cortical information flow during decision-making. Nat. Neurosci. 24, 843–850 (2021).

33. Uria, B. et al. The Spatial Memory Pipeline: a model of egocentric to allocentric understanding in mammalian brains. bioRxiv 2020.11.11.378141 (2020) doi:10.1101/2020.11.11.378141.

34. Yamins, D. L. K. et al. Performance-optimized hierarchical models predict neural responses in higher visual cortex. Proc. Natl. Acad. Sci. 111, 8619–8624 (2014).

35. Gallego, J. A. et al. Cortical population activity within a preserved neural manifold underlies multiple motor behaviors. Nat. Commun. 9, 4233 (2018).

36. Ma, W. J., Beck, J. M., Latham, P. E. & Pouget, A. Bayesian inference with probabilistic population codes. Nat. Neurosci. 9, 1432–1438 (2006).

37. Echeveste, R., Aitchison, L., Hennequin, G. & Lengyel, M. Cortical-like dynamics in recurrent circuits optimized for sampling-based probabilistic inference. Nat. Neurosci. 23, 1138–1149 (2020).

38. Remington, E. D., Narain, D., Hosseini, E. A. & Jazayeri, M. Flexible Sensorimotor Computations through Rapid Reconfiguration of Cortical Dynamics. Neuron 98, 1005–1019.e5 (2018).

39. Lu, K., Grover, A., Abbeel, P. & Mordatch, I. Pretrained Transformers as Universal Computation Engines. 210305247 Cs (2021).

40. Kirkpatrick, J. et al. Overcoming catastrophic forgetting in neural networks. Proc. Natl. Acad. Sci. 114, 3521–3526 (2017).

41. Rigotti, M. et al. The importance of mixed selectivity in complex cognitive tasks. Nature 497, 585–590 (2013).

42. Fusi, S., Miller, E. K. & Rigotti, M. Why neurons mix: high dimensionality for higher cognition. Curr. Opin. Neurobiol. 37, 66–74 (2016).

43. Stachenfeld, K. L., Botvinick, M. M. & Gershman, S. J. The hippocampus as a predictive map. Nat. Neurosci. 20, 1643–1653 (2017).

## Methods bibliography

1. Voigts, J., Newman, J. P., Wilson, M. A. & Harnett, M. T. An easy-to-assemble, robust, and lightweight drive implant for chronic tetrode recordings in freely moving animals. J. Neural Eng. (2020) doi:10.1088/1741-2552/ab77f9.

2. Newman, J. P. et al. Twister3: a simple and fast microwire twister. J. Neural Eng. (2020) doi:10.1088/1741-2552/ab77fa.

3. Siegle, J. H. et al. Open Ephys: an open-source, plugin-based platform for multichannel electrophysiology. J. Neural Eng. 14, 045003 (2017).

4. Newman, J. et al. jonnew/open-ephys-pcie: Release 1.0.0. (Zenodo, 2019). doi:10.5281/zenodo.3254431.

5. Lopes, G. et al. Bonsai: an event-based framework for processing and controlling data streams. Front. Neuroinformatics 9, (2015).

6. Chung, J. E. et al. A Fully Automated Approach to Spike Sorting. Neuron 95, 1381–1394.e6 (2017).

7. Newman, J. et al. jonnew/Oat: Oat Version 1.0. (Zenodo, 2017). doi:10.5281/zenodo.1098579.

8. Tibshirani, R. Regression Shrinkage and Selection Via the Lasso. J. R. Stat. Soc. Ser. B Methodol. 58, 267–288 (1996).

9. Eisen, M. B., Spellman, P. T., Brown, P. O. & Botstein, D. Cluster analysis and display of genome-wide expression patterns. Proc. Natl. Acad. Sci. 95, 14863–14868 (1998).

10. Tenenbaum, J. B., de Silva, V. & Langford, J. C. A global geometric framework for nonlinear dimensionality reduction. Science 290, 2319–2323 (2000).

11. Van Der Maaten, L., Postma, E. & Van den Herik, J. Dimensionality reduction: a comparative. J Mach Learn Res 10, 13 (2009).

12. Kingma, D. P. & Ba, J. Adam: A Method for Stochastic Optimization. 14126980 Cs (2017).

13. Thrun, S., Burgard, W. & Fox, D. Probabilistic Robotics | The MIT Press. https://mitpress.mit.edu/books/probabilistic-robotics (2005).

