## Supplementary figures for "Spatial reasoning via recurrent neural dynamics in mouse retrosplenial cortex"

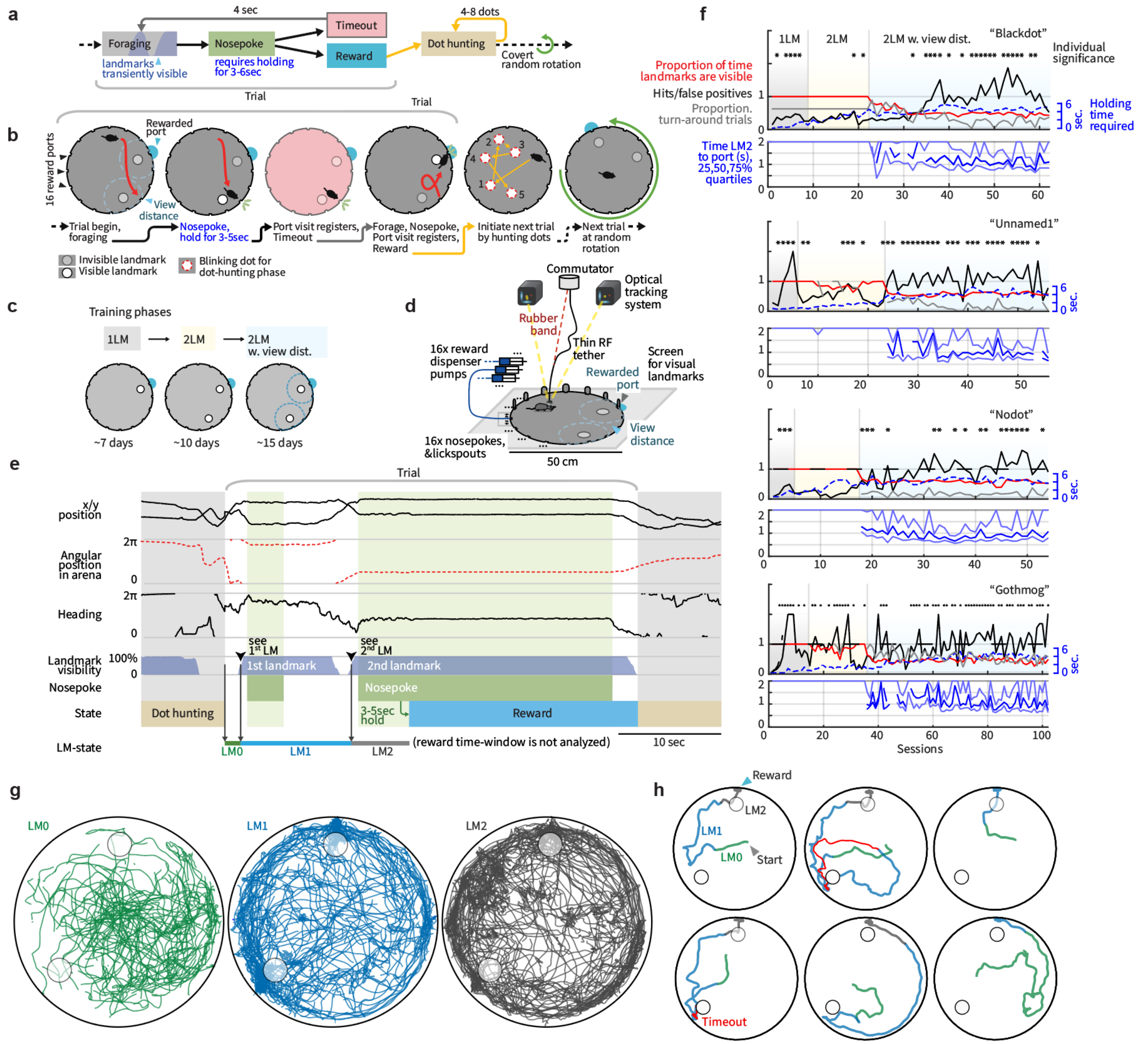


**Supplementary Figure 1.** Task structure and behavioral data. (**a**) Schematic of task structure and timing. (**b**) Example trial schematic showing all possible task states (see Methods for more details). Landmarks were formed by white dots displayed on a screen which served as the floor of the arena. They were only made visible when mice crossed a distance threshold. Only one landmark was visible at a time in the final training stage. Nose-pokes were only registered after mice held their nose in the port for a randomly chosen delay period that was randomized for each visit and not known to the mouse. Incorrect port visits resulted in timeouts that were associated with a bright background across the entire arena. After each complete trial, which results in the reward state, mice are required to complete a separate task in which they need to 'hunt' for a series of 4 to 8 randomly placed blinking dots on the arena floor. Each dot disappears as soon as the mouse reaches it, resulting either in a new random annulus, or initialization of the next trial. The next trial begins with a new random rotation of the landmarks and rewarded port. (**c**) Training phases (see Methods). Mice are trained with a single landmark first, then 2 landmarks at unlimited view distance, and finally a limited view distance. (**d**) Experimental setup for electrophysiology and real-time mouse position tracking. The arena was placed on top of a commercial flat-screen TV that was used to display visual landmarks. A motorized commutator was used to reduce tether-induced torque on the mouse, and a real-time optical tracking system was used to regulate the visibility of the landmarks and to identify when the mouse reached any of the blinking dots in the dot-hunting task. (**e**) Example excerpt of behavioral data, with state transitions. Landmark visits (black arrowheads) are defined as the point when new landmarks become visible. (**f**) Top: Training curves for all 4 mice. The three major training phases are indicated with shading (corresponding to panel c). Red: Proportion of time that a landmark is visible (remains 1.0 (100%) until view distance is introduced). Blue: Maximum reward port hold time for each session, the actual hold times are drawn from a uniform distribution. Black: proportion of hits / false positives (corresponds to rewards / timeouts, or proportion correct), for the 1st port visit in each trial. Values over 1/16 indicate that mice can distinguish the correct port amongst all ports. Values over 1 indicate that mice could reliably visit the correct port among the two ports indicated by locally ambiguous landmarks without excluding any other ports by trial and error (see main text and methods). Trials with 1st landmark visit after <20 sec are included in analysis. Grey: Proportion of trials in which mice see both landmarks, and then turn around to go back to the 1st landmark. If this proportion was 0, it would indicate that mice always visit the 2nd port after seeing it, which would on average lead to chance-level behavioral performance. For each individual session, significance of correct choice for the 1st port visit among the two indicated ports was tested with a binomial fit at the 95% level and is indicated with a star. Bottom: latency to reward after encountering the 2nd landmark in seconds. (**g**) All paths taken by the mouse in one example session, split by LM0,1,2 state (green, glue, grey). (**h**) 6 example trials from the same session plotted from the start of the trial to the reward delivery, same color scheme as in g. 2 of the trials include time-outs (red).

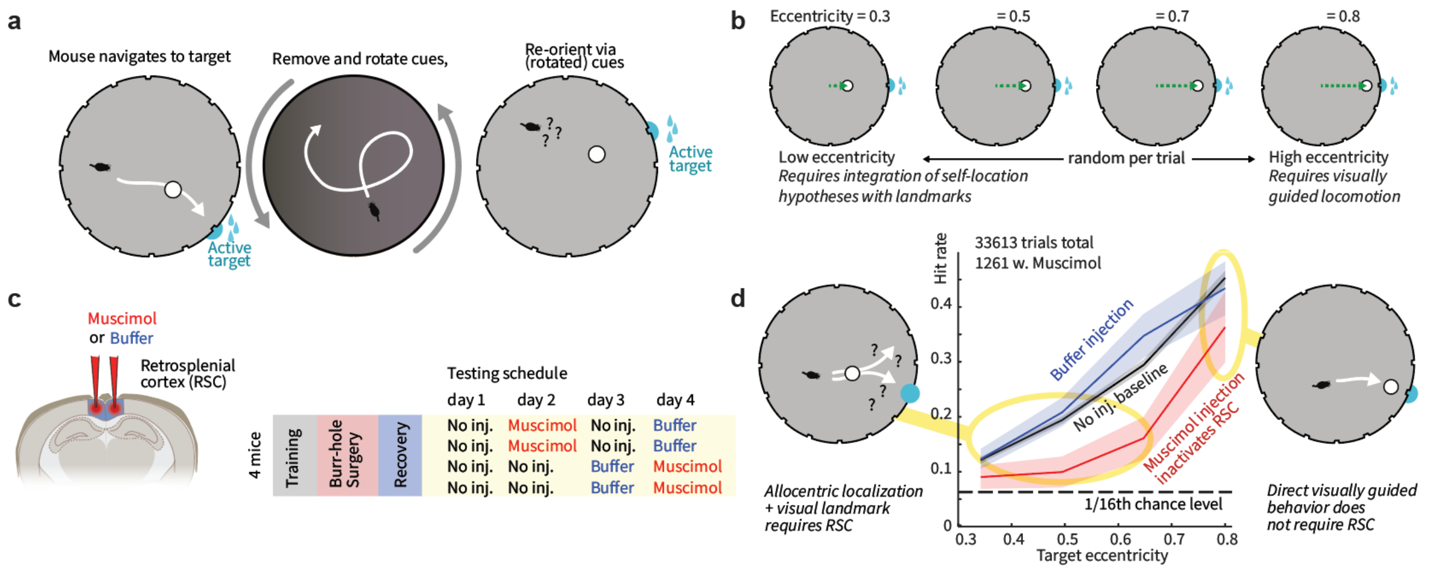


**Supplementary Figure 2**. Retrosplenial cortex is required for integrating egocentric sensory information and allocentric spatial hypotheses, but not for visually guided navigation. To causally test the role of RSC in relating spatial hypotheses to sensory data, we used a parametric allocentric/egocentric task using the same apparatus as in the main experiment and pharmacologically inactivated RSC. (**a**) Schematic of task structure. Water restricted mice had to visit the port closest to a single visual landmark for a water reward. Visits to any other port resulted in a time-out, but allowed the mice to self-correct. As in the main experiment, the landmark and rewarded port were rotated randomly after each trial, forcing mice to use only the visual landmark. (**b**) To make the task reliant on allocentric hypotheses, we randomly varied the eccentricity of the landmark at the beginning of each trial. Trials with low eccentricity (left) required the mouse to find the arena center (though path integration) and then extrapolate a straight path through the landmark to the correct rewarded port. Alternatively, mice might triangulate which port is the closest to the landmark from the periphery. These strategies all require integration of self-location hypotheses with visual landmark information. Trials with high eccentricity (right) required merely walking to the port closest to the landmark. This design allowed us to test the role of RSC in the integration of location hypotheses with egocentric visual landmark information while simultaneously determining whether simpler visually-guided navigation was also affected. (**c**) RSC was either 1) transiently inactivated with Muscimol, 2) sham injected with cortex buffer, or 3) not injected (see methods). Each mouse was tested in both groups, with balanced ordering. (**d**) Task performance (95% confidence intervals for hit rate on 1$^{st}$ port visits per trial, via binomial bootstrap). Mice always performed above full chance level (1/16$^{th}$, assuming they cannot make use of the landmark). Performance was selectively reduced by RSC inactivation for low eccentricity conditions where integration of location hypotheses and visual landmarks was required. Performance in the visually guided condition was only minimally affected.

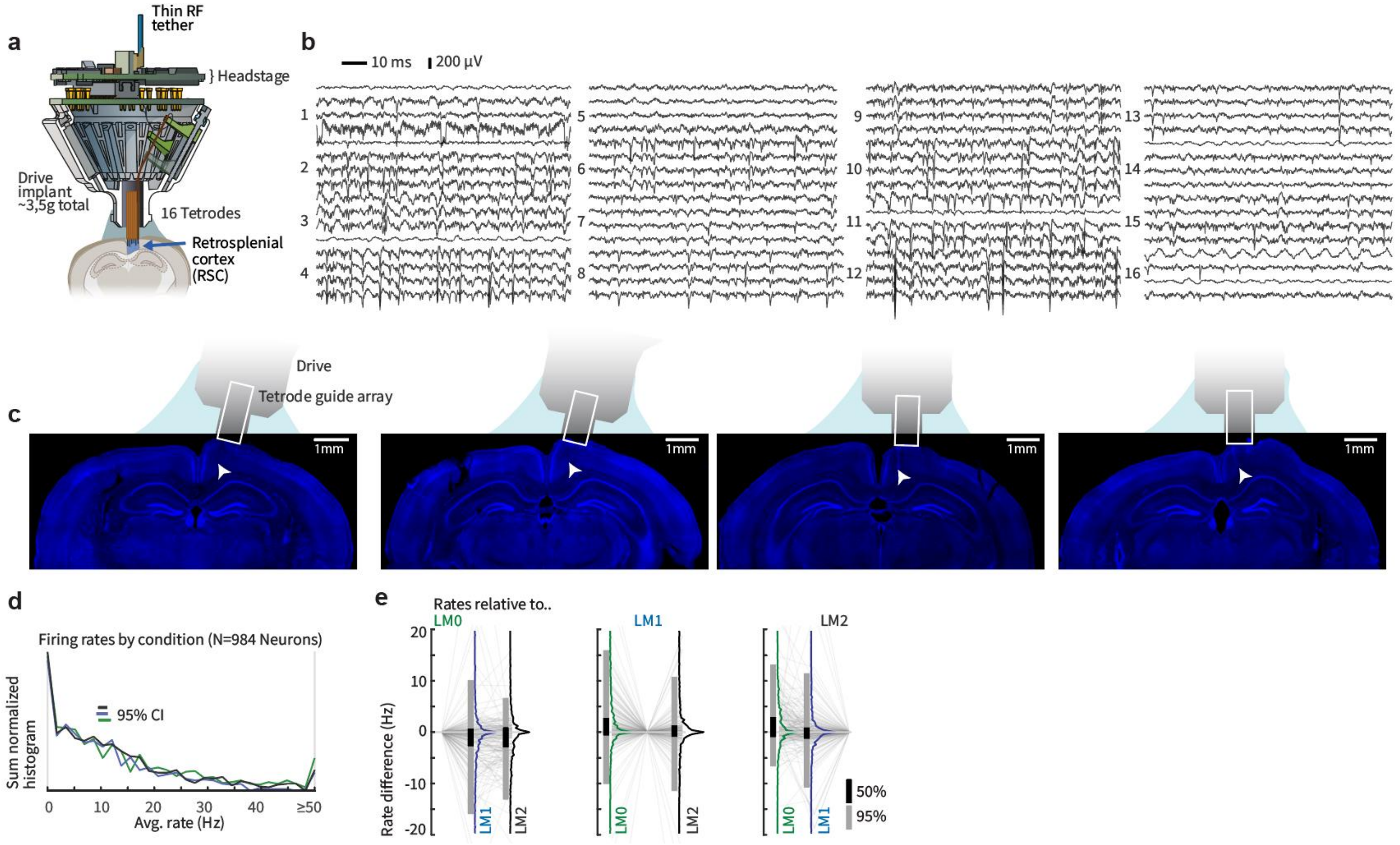


**Supplementary Figure 3.** Extracellular recording in mouse retrosplenial cortex. (**a**) Tetrode drive[1] implants targeting mouse retrosplenial cortex (RSC). See Methods for details. (**b**) Example band-passed (100Hz-5kHz) raw voltage traces from 16 tetrodes. (**c**) Verification of drive implant locations in RSC via histology in all 4 mice. White arrowheads indicate electrolytic lesion sites. (**d**) Histograms of mean firing rates of all 984 neurons across LM0 (green), LM1 (blue), and LM2 (black) conditions. Overall rates did not shift significantly across these states. (**e**) Relative per-neuron changes in firing rates across conditions. Despite the lack of a population-wide shift in average rates, the firing rates of individual cells varied significantly across conditions with heterogeneous patterns of rates. Each grouping shows rates per cell, relative to the rate in LM0 (left) LM1 (middle), and LM2 (right) as individual rates (grey lines and histograms). Bar graphs show the 50% and 95% quantiles.

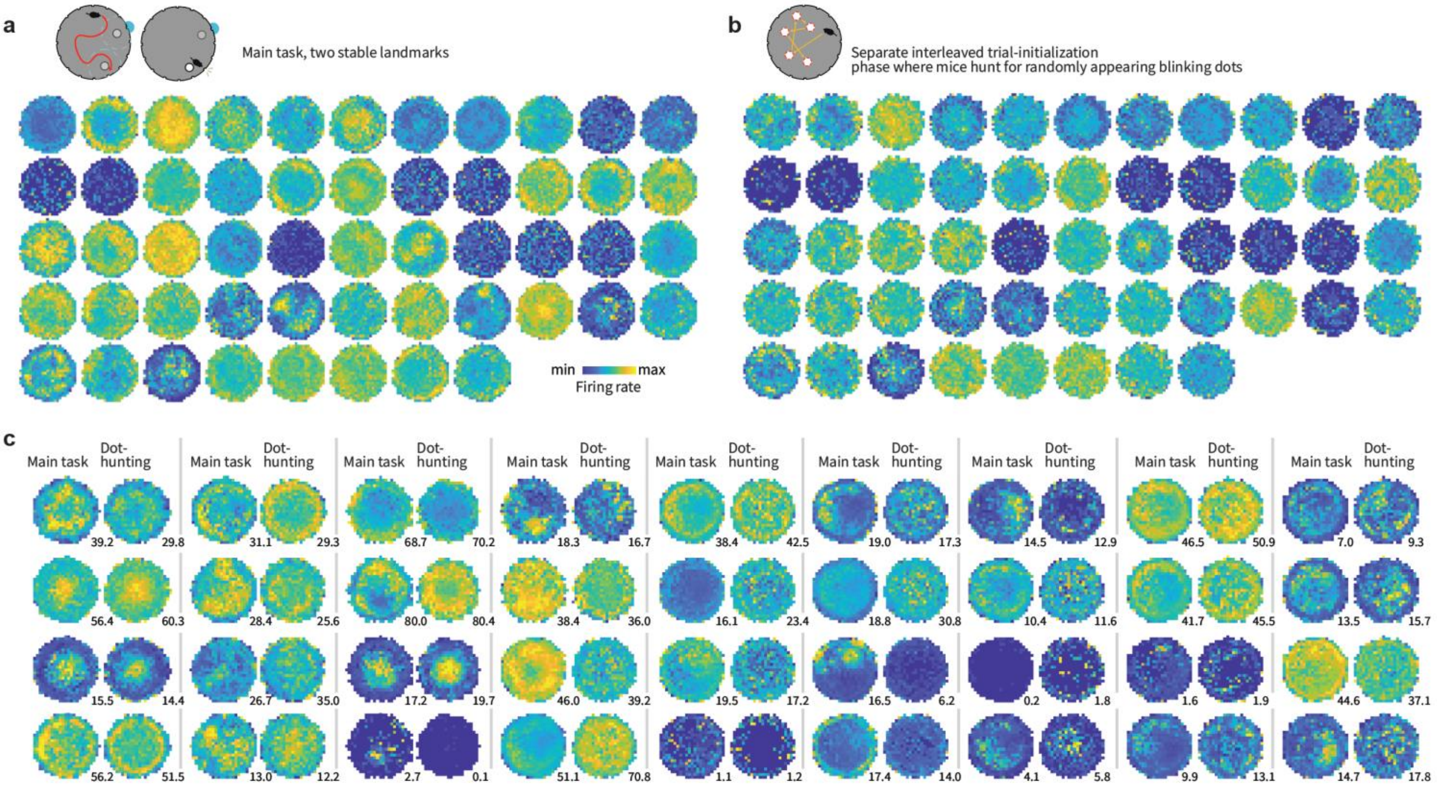


**Supplementary Figure 4.** Example spatial firing rate profiles of neurons in mouse retrosplenial cortex. (**a**) Spatial firing rate profiles for all neurons from one example session (52 total), from the main task phase. Profiles were computed in 25x25 bins, and individually normalized to their 99$^{th}$ percentile. (**b**) same as panel a, but from the separate trial initialization task (‘dot-hunting’) in which mice had to hunt for a series of blinking dots that appeared in random positions. (**c**) 36 example neurons from multiple sessions and animals, chosen to represent the broad range of tuning profiles. For each neuron, the main task tuning and the ‘dot-hunting’ are plotted together on the same brightness scale, normalized to their total maximal rate. In the dot-hunting task there is no conserved radial tuning due to the absence of consistent landmarks, however some cells retain angular spatial tuning due to olfactory cues in the arena. Tuning to eccentricity (distance to arena wall or center) is maintained across task phases in many neurons. Small numbers indicate maximum firing rates in Hz for each plot (color scale is same across the pairs).

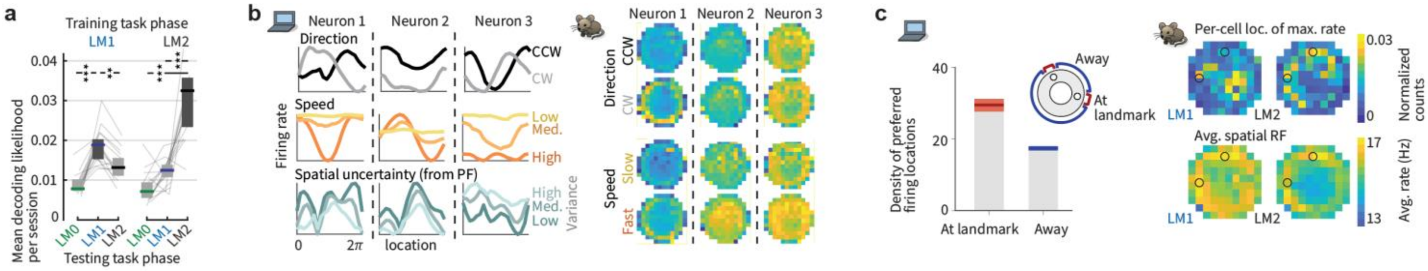

**Supplementary Figure 5.** The spatial code in RSC changes with hypothesis states, ANN and RSC neurons employ conjunctive codes, and preferentially represent landmark / reward locations. (**a**) Location decoders (neural network, cross-validated per trial) do not generalize across LM states, and LM1 carries less spatial information than LM2. Performance is measured by prediction likelihood in a 10x10 grid, shading: 95% CIs across sessions. See Fig.1f for test via spatial RF differences. (**b**) Left: Example ANN neuron tuning curves (from LM2) split by travel direction, speed, or location uncertainty (corresponding to LM0,1,2 states, derived from particle filter), showing conjunctive coding. Right: Three RSC example cells showing conjunctive coding of location vs. speed, and direction (Fig.1d shows task phase vs. location). (**c**) Left: ANN neurons and, Right: RSC cells (N=984 neurons) preferentially fire at landmark locations. Top: distribution of locations where RSC cells fire most. Bottom: total average rates, split by LM1 and LM2.

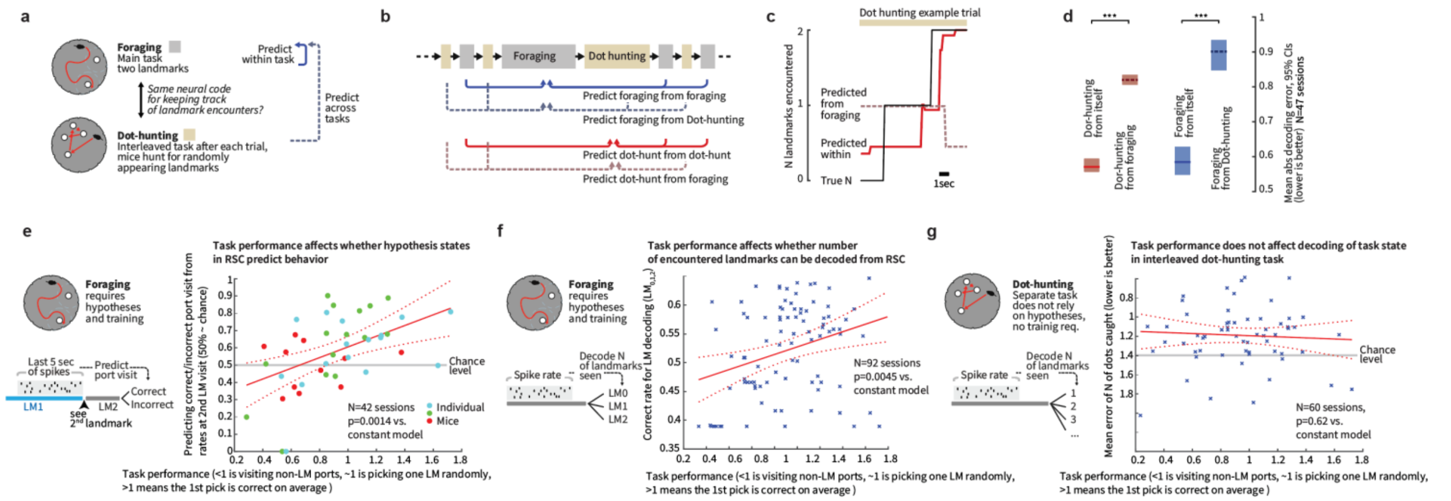

**Supplementary Figure 6.** Hypothesis encoding in RSC is task-specific and is a function of task learning. (**a**) Foraging and dot-hunting tasks are interleaved, allowing us to compare how the same neural population represents hypotheses. (**b**) We predict the number of encountered landmarks either within condition (e.g. foraging from foraging, each time using one trial as test, fitting to all others). Only the first 2 landmarks were predicted to allow use of the same classifier across both despite the higher number of landmarks in the dot-hunting task. Train and test sets were split by trial. Decoding was performed with a regression tree on low-pass filtered firing rates. Performance was quantified as mean error on the N of landmarks. (**c**) Example dot-hunting trial, the performance from using the foraging predictor is lower. (**d**) Summary stats from all sessions. The prediction is significantly better when using training data from the same category than when using the neural code from the other; e.g. dot-hunting to predict the foraging (P= ~0 / ~0 within vs. across categories for predicting dot-hunting and foraging LM state), showing that hypothesis coding is task-specific. (**e**) To test whether hypothesis encoding is a specific function of task learning or a general feature of RSC, we examined whether coding persisted in case when mice performed the task, but were not yet performing well. We first examined the ability to predict correct vs. incorrect port choice (same as in Fig. 4) as a function of per-session task performance. We analyzed data from sessions from the entire training period where the 2 landmarks were used, with at least 5 correct and 5 incorrect choices, yielding N=42 sessions total. On average we analyzed ~15-30 port visits per session (number of trials was unaffected by behavioral performance: CI of slope = [-7.7, 2.7], p = 0.33). Predictions were made as before with a test/train split on balanced hit/miss data with a regression tree. Prediction performance was at chance level (~47%, P=0.81 vs. chance) for low performance sessions (total correct choice ratio of 0.8 or lower), and the same as in our initial analysis (Fig. 4) for sessions with high mouse performance (~66%, P=0.00096 vs. chance). Overall, prediction performance was significantly correlated with task performance (P=0.0014 vs. constant model). Individual mice are indicated with colored markers. (**f**) We also analyzed the more general decoding of landmark encounter count (same as Fig. 1) in all of the 92 sessions with 2 landmarks, and also found a significant correlation (p=0.0045 vs. constant model), showing that hypothesis encoding throughout the task is driven by task learning. (**g**) As a control experiment, we tested whether decoding the number of landmarks encountered in the interleaved dot-hunting task might also be affected by task performance, if for instance the neural encoding and performance was a function of general spatial learning, habituation to the arena, motivation, etc., and we found this correlation to be flat (P=.6, CI for slope = [-0.17, 0.29]). We conclude that the encoding of hypothesis state is task-specific and a function of the mouse performing the task.

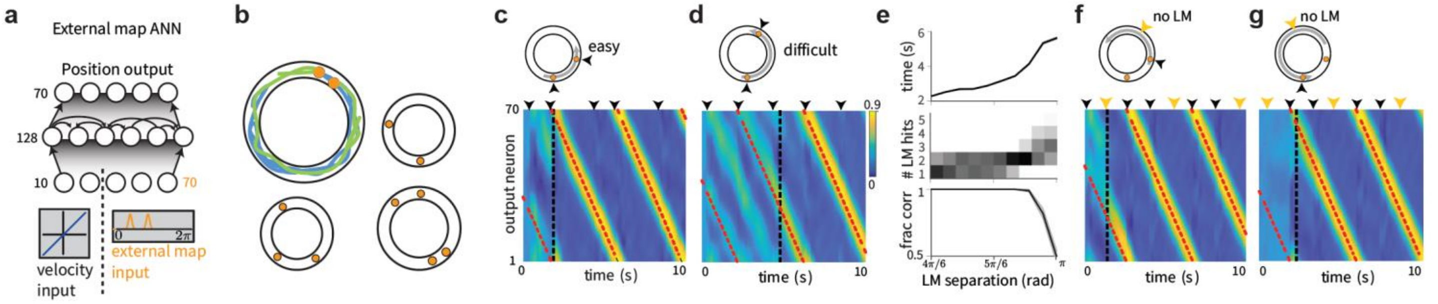


**Supplementary Figure 7.** Architecture and trajectories for ANN with external map input. (**a**) Structure of the recurrent network. Input neurons encoded noisy velocity input with linear tuning curves (similar to speed cells in the entorhinal cortex[2]), and landmark (LM) information. In the standard setup (which we refer to as "external map"), the LM input signaled the global configuration of LMs (map). If there are K LMs in the environment (all assumed to be perceptually indistinguishable), then whenever the animal encounters one of the LMs, the input provides a simultaneous encoding of all K LM locations using spatially tuned input cells. Thus, the input encodes the map of the environment but does not disambiguate locations within it. This input can be thought of as originating from a distinct brain area that identifies the current environment and provides the network with its map. (**b**) Trajectories varied randomly and continuously in speed and direction. There were 2-4 LMs at random locations. (**c**) Activity of output neurons ordered by preferred location as a function of time in an easy trial with two nearby LMs and a constant velocity trajectory. Black arrows: time of LM encounters. Thick black dashed line: time of disambiguation of location estimate in output layer. Thin red dashed line: true location. The network's decision on when to collapse its estimate is flexible, and dynamically adapts the decision time to task difficulty: When the task is harder because of the configuration of LMs (the task becomes harder as the two LMs approach a 180 degree separation because of velocity noise and the resulting imprecision in estimating distances; the task is impossible at 180 degree because of symmetry), the network keeps alive multiple hypotheses about its states across more LM encounters until it is able to reach an accurate decision. Panels c,d,f,g show example trials from experiment configuration 4 (See Methods) with different values of landmark separation parametrized by α. (**d**) Same as c, but in a difficult trial with two LMs almost opposite of each other. (**e**) Top: The ANN took longer to disambiguate its location in harder task configurations: average time until location disambiguation as a function of LM separation (Standard error bars are narrower than line width). Middle: Distribution of the number of LM encounters until the network disambiguates location, as a function of LM separation. Bottom: Fraction of trials in which the network location estimate is closer to the correct than the alternative LM location at the last LM encounter, as a function of LM separation. Data from 10000 trials in experiment configuration 4, 1000 for each of the 10 equally spaced values of α. The performance of the ANN (Fig. 2 main text) can be compared to the much poorer performance achieved by a strategy of path integration to update a single location estimate with LM-based resets (to the coordinates of the landmark that is nearest the current path-integrated estimate), Fig. 2b (black versus gray). The latter strategy is equivalent to existing continuous attractor integration models[3,4] combined with a LM- or border based resetting mechanism[5–8], which to our knowledge is as far as models of brain localization circuits have gone in combining internal velocity-based estimates with external spatial cues. The present network goes beyond a simple resetting strategy, matching the performance of a sequential probabilistic estimator – the particle filter (PF) – which updates samples from a multi-peaked probability distribution over possible locations over time and is asymptotically Bayes-optimal (M = 1000 particles versus N = 128 neurons in network; Fig. 2b, lavender (PF) and green (enhanced PF)). Notably, the network matches PF performance without using stochastic or sampling-based representations, which have been proposed as possible neural mechanisms for probabilistic computation[9,10]. (**f**) Similar to c, but in a trial where the network disambiguates its location before the second LM encounter. Yellow arrows mark times of LM interactions if the alternative location hypothesis had been correct. Disambiguation occurs shortly after the absence of a LM encounter at the first yellow arrow. (**G**) Similar to f, but in a trial where disambiguation occurs at the first LM location, since no LM has been encountered at the time denoted by the first gray arrow.

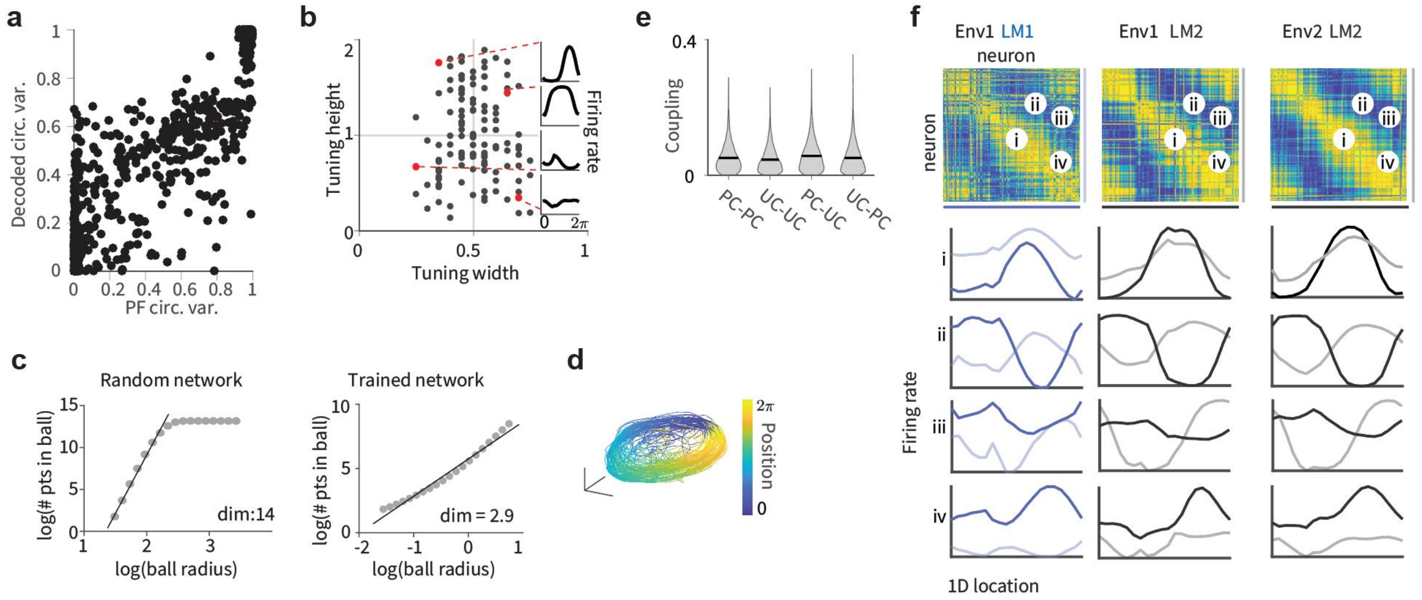


**Supplementary Figure 8.** Population statistics for ANN with external map input. (**a**) Scatter plot of enhanced particle filter (ePF) circular variance vs. estimate decoded from hidden layer of the network. 4000 trials from experiment configuration 1 were used to train a linear decoder on the posterior circular variance of the ePF from the activity of the hidden units and performance was evaluated on 1000 test trials. (**b**) Scatter plot of widths and heights of ANN tuning curves after the 2nd LM encounter. Insets: example tuning curves corresponding to red dots. Unlike hand-designed continuous attractor networks, where neurons typically display homogeneous tuning across cells[4,11,12], our model reproduces the heterogeneity observed in hippocampus and associated cortical areas. Tuning curves are from LM2 using 1000 trials from experiment configuration 2 using 20 location bins. Tuning height specifies the difference between the tuning curve maximum and minimum, and tuning width denotes the fraction of the tuning curve above the mean of maximum and minimum. (**c**) State-space activity of ANN is approximately 3-dimensional. Even when summed across all environments and random trajectories, the states still occupy a very low-dimensional subspace of the full state space, quantified by the correlation dimension as d ≈ 3 (left, see Methods). This measure typically overestimates manifold dimension[13], and serves as an upper bound on the true manifold dimension. As a control, the method yields a much larger dimension (d = 14) on the same network architecture with large random recurrent weights (right); thus, the low-dimensional dynamics are an emergent property of the network when it is trained on the navigation task. Data from 5000 trial, recurrent weights were sampled i.i.d. from a uniform distribution $W_{h,ij} \sim U([-1, 1])$, then fixed across trials. The initial hidden state across trials was sampled from $h_{t=0,i} \sim U([-1, 1])$. Data from 5000 trials from experiment configuration 1. (**d**) In the LM2 state, position on the rate-space attractor corresponds to location in the maze. State-space trajectories after second LM encounter for random trajectories. Color corresponds to true location (plot shows 100 trials). (**e**) The distribution of recurrent weights shows that groups of neurons with strong or weak location tuning or selectivity have similar patterns and strengths of connectivity within and between groups: distribution of absolute connection strength between and across location-sensitive "place cells" (PCs) and location-insensitive "unselective cells" (UCs) in the ANN. The black line denotes the mean; s.e.m. is smaller than the linewidth. The result is consistent with data suggesting that place cells and non-place cells do not form distinct sub-networks, but are part of a system that collectively encodes more than just place information[14]. Location tuning curves were determined after the second landmark encounter using 5000 trials from distribution 1 and using 20 location bins. The resulting tuning curves were shifted to have minimum value 0 and normalized to sum to one. The location entropy of each neuron was defined to be the entropy of the normalized location tuning curve. Neurons were split in two equal sets according to their location entropy, where neurons with low entropy were defined as "place cells" (PCs) and neurons with high entropy were defined as "non-place cells" (UCs). Between and across PCs and UCs absolute connection strength was calculated as the absolute value of the recurrent weight between non-identical pairs. (**f**) Pairwise correlation structure[15] is maintained across LM[1,2] states and environments. Corresponds to Fig. 3a. Top: Correlations in spatial tuning between pairs of cells in one environment after the 1st LM encounter / LM1 (left), after the 2nd encounter / LM2, and in a separate environment in LM2 (right). The neurons are ordered according to their preferred locations in environment 1. Bottom: Example tuning curve pairs (normalized amplitude) corresponding to the indicated locations i-iv. Data from experiment configuration 1.

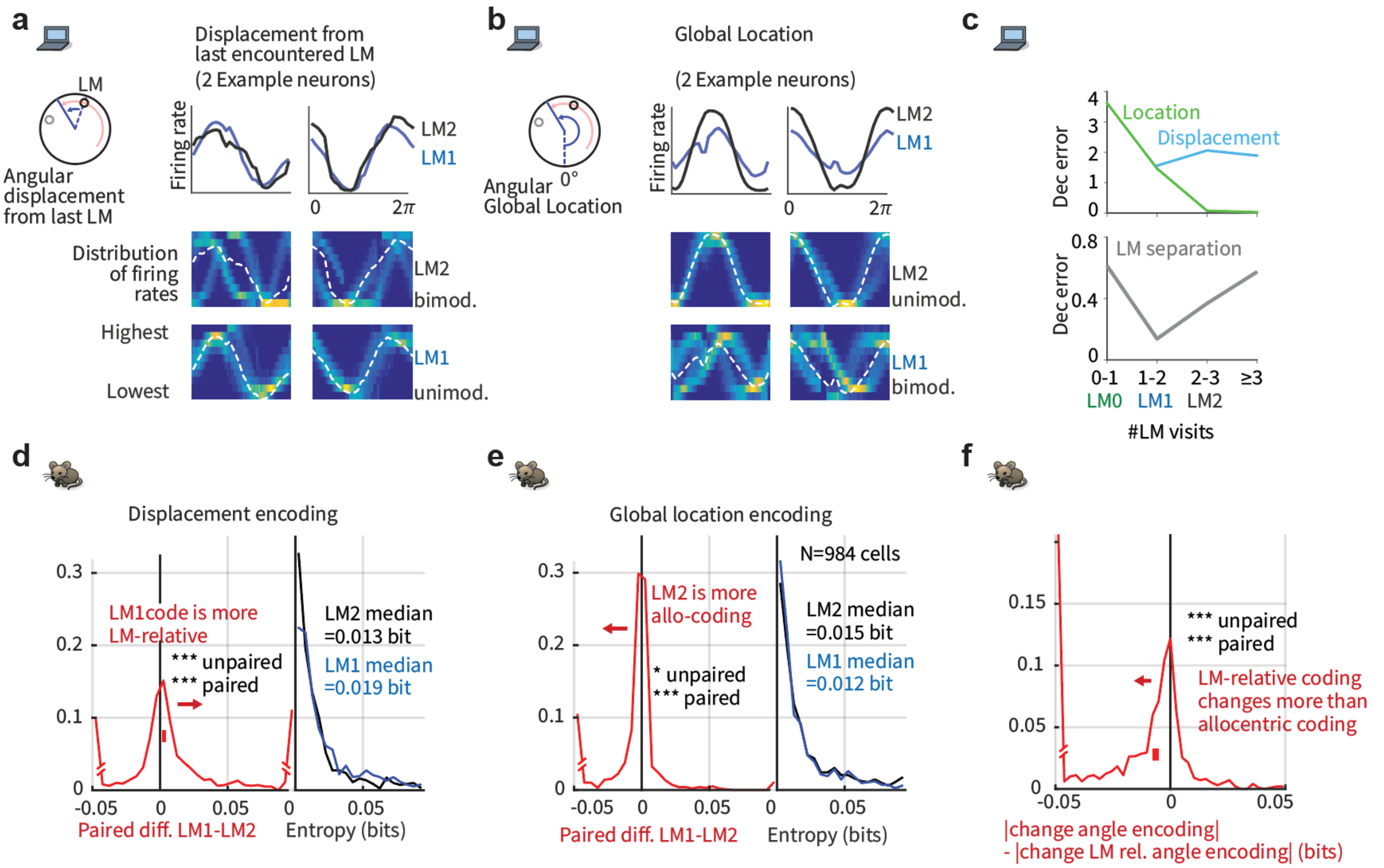


**Supplementary Figure 9.** ANN and RSC coding transitions dynamically from an egocentric landmark-relative to an allocentric global reference frame based on phase in trial. (**a**) Top: Tuning curves (mean rate) for displacement from last encountered LM for LM1 and LM2 states in ANN. Bottom: Same data, but distribution of firing rates. The network discovers that displacement from the last LM encounter in the LM1 period is a key latent variable, and its encoding is an emergent property. Intriguingly, a similar displacement-to-location coding switch has been observed in mouse CA1[16], suggesting that the empirically observed switch may be related to the brain performing spatial reasoning to disambiguate between multiple location hypotheses. (**b**) Same as panel a but for global location, ANN neurons became more tuned to global location rather than LM-relative information after encountering the 2$^{nd}$ landmark. (**c**) Decoding of location, displacement, and separation between landmarks from the ANN in a 2-landmark environment by a linear decoder that remains fixed across trials and environments. Top: Squared population decoding error of location (green) and displacement (blue), as a function of the number of encountered LMs. As suggested by the well-tuned activity of ANN neurons, location can be linearly decoded in the LM2 state. Displacement can be best decoded in the LM1 state. Bottom: Square decoding error of distance between LMs, as a function of the number of encountered LMs. The representation is particularly accurate around the time just before and after the first LM encounter, when location disambiguation takes place. Top: Performance was evaluated on 1000 trials from experiment configuration 2. For location, the decoder corresponded to the network location estimate. For displacement, the linear decoder was trained on 4000 separate trials. Bottom: experiment configuration 1 with 4000 trials to train the linear decoder and 1000 trials to evaluate it. Thus, the network's encoding of these three critical variables is dynamic and tied to the different computational imperatives at each stage: displacement and LM separation are not explicit inputs but the network estimates these and represents them in a decodable way at LM1, the critical time when this information is essential to the computation. After LM2, the network decodability of LM separation drops, as it is no longer essential. (**d**) Neurons in RSC also became less well tuned to relative displacements from landmarks in LM2 relative to LM1: histogram across all RSC neurons of entropy of tuning curve for angular displacement from last seen landmark in RSC. Black: for LM2 state, Blue: for LM1 state. Red: histogram of pairwise differences. For this analysis, angular firing rate distributions were analyzed relative to either the global reference frame or the last seen landmark. (**e**) Same as D, but for global location. (**f**) The absolute change in landmark-relative displacement coding (d) is larger than that of the allocentric location tuning (e), suggesting that the latter is less affected by task state.

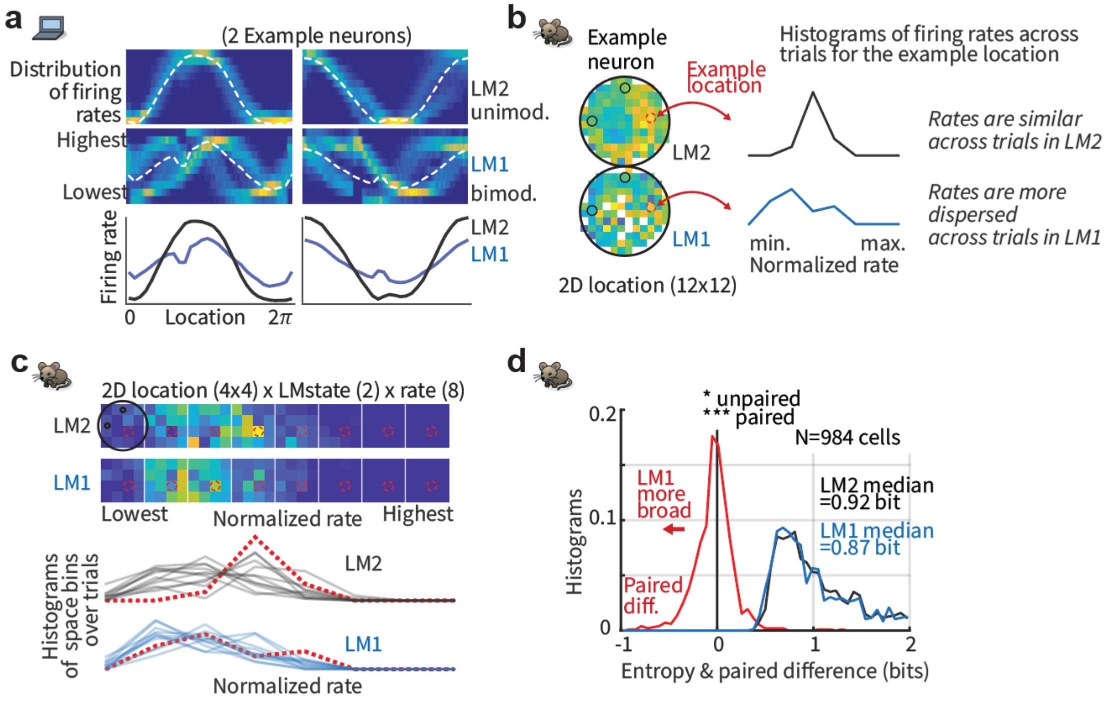


**Supplementary Figure 10.** In addition to explicitly encoding number of visited landmarks, RSC and the ANN exhibit higher trial-to-trial variability in partial information states. (**a**) Bottom: Mean spatial activity profile of 2 example ANN neurons for LM1 and LM2. Average tuning is higher for the LM2 state. Top: same data as histograms, showing that the less well-tuned LM1 state corresponds to a bimodal rate distribution (rates are high in some trials, low in others) that transitions to a unimodal distribution once the 2$^{nd}$ landmark has been identified in LM2. Data are from experiment configuration 2 (See Methods, section 'Overview over experiment configurations used with ANNs'). Tuning curves were calculated using 20 bins of location/displacements and normalized individually for each neuron. The first time step in each trial and time steps with non-zero landmark input were excluded from the analysis. For histograms, each condition was binned in 100 column bins and neuron rates in 10 row bins. Histograms were normalized to equal sum per column. (**b**) Similarly, RSC rates are more dispersed per location in LM1. Schematic of analysis: firing rates were low pass filtered at 0.5 Hz, and for each location, the distribution of rates was computed in 8 bins, between the lowest and highest rate of that cell. (**c**) Example analysis for one cell. Top: Rate distribution resolved by 2D-location (4x4 bins) for example RSC neuron. Bottom: the resulting 16 histograms for LM1 and LM2 each, red dotted example histograms correspond to indicated example location (red dotted circles). (**d**) Summary statistics showing a more dispersed rate distribution per location in LM1.

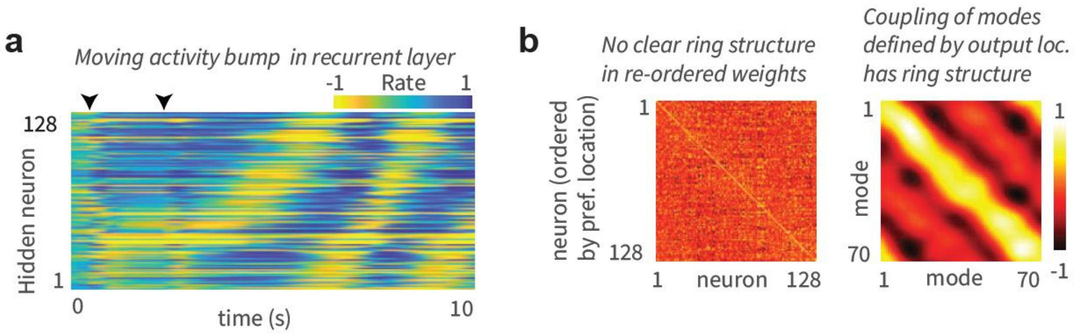


**Supplementary Figure 11.** ANN with external map input implements a circular attractor structure. (**a**) Hidden layer activity arranged by preferred location in an example trial shows a bump of activity that moves coherently. Black arrows: first two LM encounters. Preferred location was determined after the second landmark encounter using 5000 trials from experiment configuration 1. (**b**) Left: Recurrent weight matrix arranged by preferred location of neurons (determined after the second landmark encounter using 5000 trials from experiment configuration 1) indicates no apparent ring structure, despite apparent bump of activity that moves with velocity inputs (panel a). Right: However, recurrent coupling of modes defined by output weights (defined by $W_{out}\ W_{rec}W_{out}^{T}$, where $W_{rec}$ are the recurrent weights and $W_{out}$ are the output weights) has a clear band structure. Connections between appropriate neural mixtures in the hidden layer – defined by the output projection of the neurons – therefore exhibit a circulant structure, but the actual recurrent weights do not, even after sorting neurons according to their preferred locations. The ANN thus implements a generalization of hand-wired attractor networks, in which the integration of velocity inputs by the recurrent weights occurs in a basis shuffled by an arbitrary linear transformation. Given these results, one cannot expect a connectomic reconstruction of a recurrent circuit to display an ordered matrix structure even when the dynamics are low-dimensional, without taking into account the output projection. Because trials in the mouse experiments typically ended almost immediately when the mouse had seen both landmarks (See Supplementary Fig. 1f for a quantification), we did not quantify the topology of the neural dynamics in RSC.

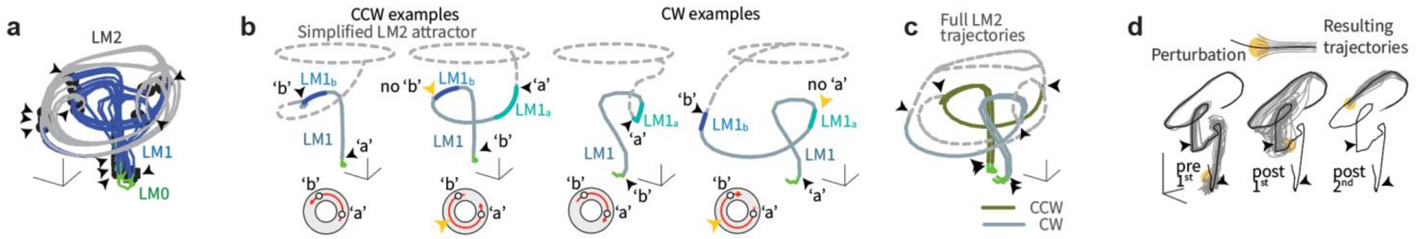


**Supplementary Figure 12.** Low-dimensional state-space dynamics in the ANN with external map input suggests novel form of probabilistic encoding. (**a**) Visualization of the full state-space dynamics of the hidden layer population, projected onto the three largest principal components, for constant-velocity trajectories. ANN hidden layer activity was low-dimensional: Fig. 3a shows data on low-dimensional dynamics, evident in maintained pairwise correlations, and Figs. 3d and Supplementary Fig. 8c show correlation dimension. Trajectories are shown from the beginning of the trials; arrows indicate LM encounter locations, black squares: first LM encounter; black circles: second LM encounter; line colors denote trajectory stage: LM0 (green), LM1 (blue), andLM2 (grey). Data in a-c is from 1000 trials from experiment configuration 3 (see Methods); sensory noise was set to zero. Trajectory starting points were selected to be a fixed distance before the first landmark. The intermediate ring (LM1) corresponds to times at which the output neurons represent multiple hypotheses, whereas the final location-coding ring (LM2), well-separated from the multiple hypothesis coding ring, corresponds to the period during which the output estimate has collapsed to a single hypothesis. In other words, the network internally encodes single-location hypothesis states separably from multi-location hypothesis states, as we find in RSC (Fig. 1), and transitions smoothly between them, a novel form of encoding of probability distributions that appears distinct from previously suggested forms of probabilistic representation[9,10]. (**b**) ANN trial trajectory examples, (corresponding to Fig. 2e). Divergence of trajectories for two paths that are idiothetically identical until after the second LM encounter. ‘a’ and ‘b’ denote identities of locally ambiguous identical landmarks. Disambiguation occurs at the second LM encounter, or by encountering locations where a LM would be expected in the opposite identity assignments. See insets for geometry of trajectories and LM locations. LM2 state has been simplified in these plots. (**c**) All four trajectories from panel b plotted simultaneously, and with full corresponding LM2 state. (**d**) The low-dimensional state-space manifold is stable, attracting perturbed states back to it, which suggests that the network dynamics follow a low-dimensional continuous attractor and the network’s computations are robust to most types of noise. Relaxations in state space after perturbations before the first (left), between first and second (middle), and after the second (right) LM encounter. For the base trial, a trial with two landmarks and random trajectory was chosen. The first and second landmark encounter in this base trial is at time t = 2s and t = 4.6s respectively. At time t = 1s (left), t = 4s (middle), and t = 7s (right) a multiplicative perturbation of size 50% was introduced at the hidden layer.

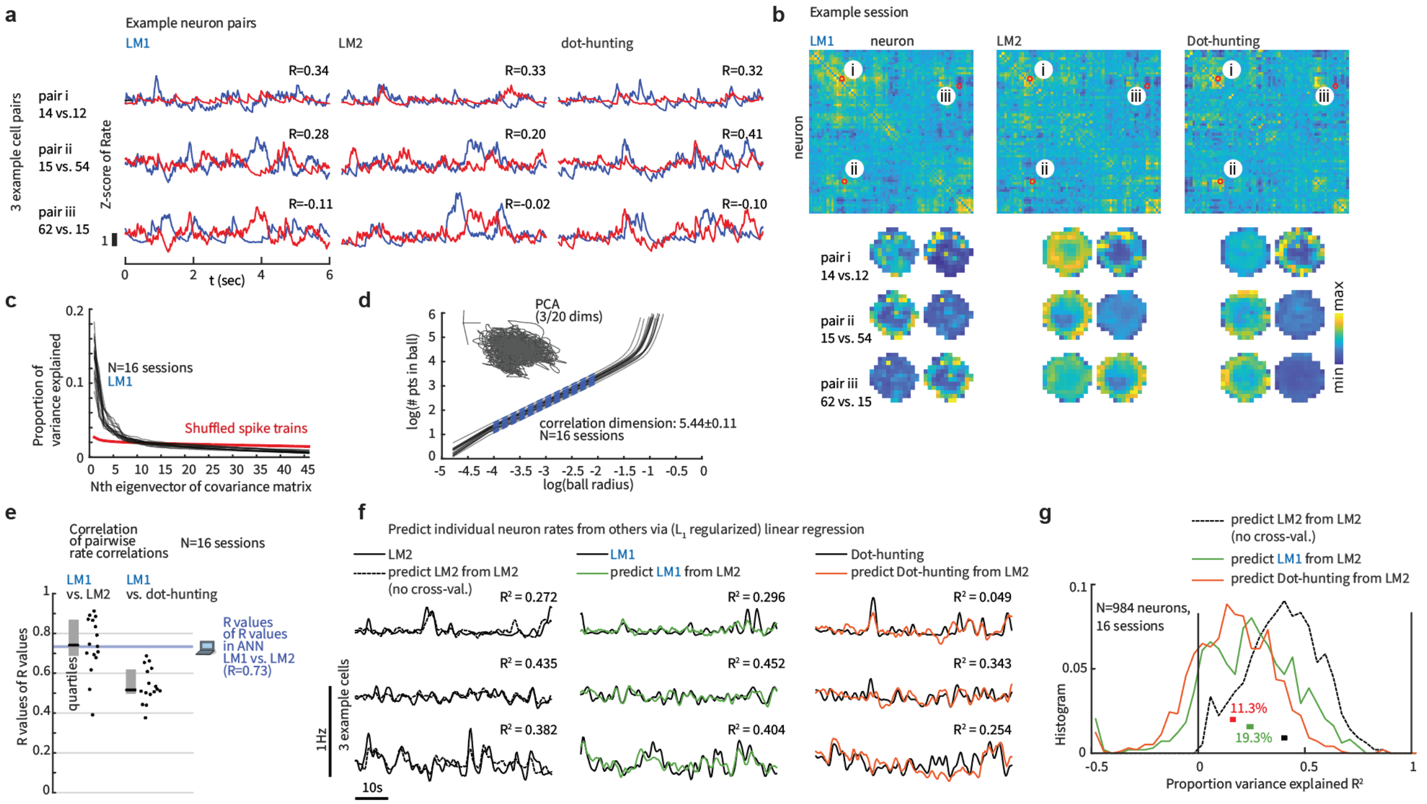


**Supplementary Figure 13.** Pairwise rate correlation structure in RSC is maintained across LM1 and LM2 states. Low-dimensional population structure can be probed by pairwise neural relationships[15]: correlations or offsets in spatial tuning between cell pairs should be preserved across environments if the dynamics across environments is low-dimensional. (**a**) Example spike rates (6 sec window, low-passed at 1Hz using a single-pole Butterworth filter) for 3 RSC neuron pairs from one example session. R values for each pair were computed across the LM1 and LM2 condition, as well as in the task-initialization phase where mice had to hunt blinking dots (Supplementary Figs. 1a, 4). The latter provides a control condition where no landmark based navigation was required and mice instead had to walk to randomly appearing targets. (**b**) Top: pairwise correlation matrices for LM1,2 and dot-hunting conditions. Example pairs are highlighted (i,ii,iii). Bottom: spatial firing rate profiles for example pairs. Same analysis as in Fig. 3a. (**c**) RSC activity is globally low-dimensional. Proportion of variance of low-pass filtered (0.5Hz) firing rates explained by first 45 principal components from the LM1 states. Proportion of variance explained (black, 16 sessions) drops to below that of shuffled spike trains (red) after the 6-10$^{th}$ principal component. (**d**) Correlation dimension in RSC is also low (same analysis as for the ANN in Supplementary Fig. 8c). This measure typically overestimates manifold dimension[13], and thus serves as an upper bound on the true manifold dimension. (**e**) Summary statistics for correlation of correlations (panel b shows one example session). Median of R value of R values for LM1 vs. LM2 = 0.74 (corresponding R in ANN = 0.73), for LM1 vs. dot-hunting = 0.51. (**f**) Rates of individual RSC neurons can be predicted from other neurons with linear regression. In the LM2 to LM2 condition (black), the linear fit is computed for one held-out neuron's rate from other concurrent rates, and the same regression weights are then used to predict rates during LM1 (green) and dot-hunting (red) time periods. True rates of predicted neurons are plotted as solid black lines. (**g**) Summary statistics for the linear regression. Histograms show the proportion of explained variance for all 984 neurons, split by condition. In the LM2 to LM2 condition, the fit is computed from other concurrent rates (40.5% variance explained). In the two other conditions, the regression weights are fit in LM2 and held fixed. The sequential, non-interleaved nature of this train/test split across task phases means that any consistent firing rate drifts across the conditions will lead to poor predictions, and consequently, a small number of neurons exhibit negative $R^2$ values indicating a fit that is worse than an average rate model (11.3% for LM1, 19.3% for dot-hunting). However, even under these conditions, 24.3% of variance (median across neurons) can be explained despite significant changes in spatial receptive fields (predict LM1 with LM2 weights) and even for a different task, with 16.2% when predicting dot-hunting activity from LM2 weights.

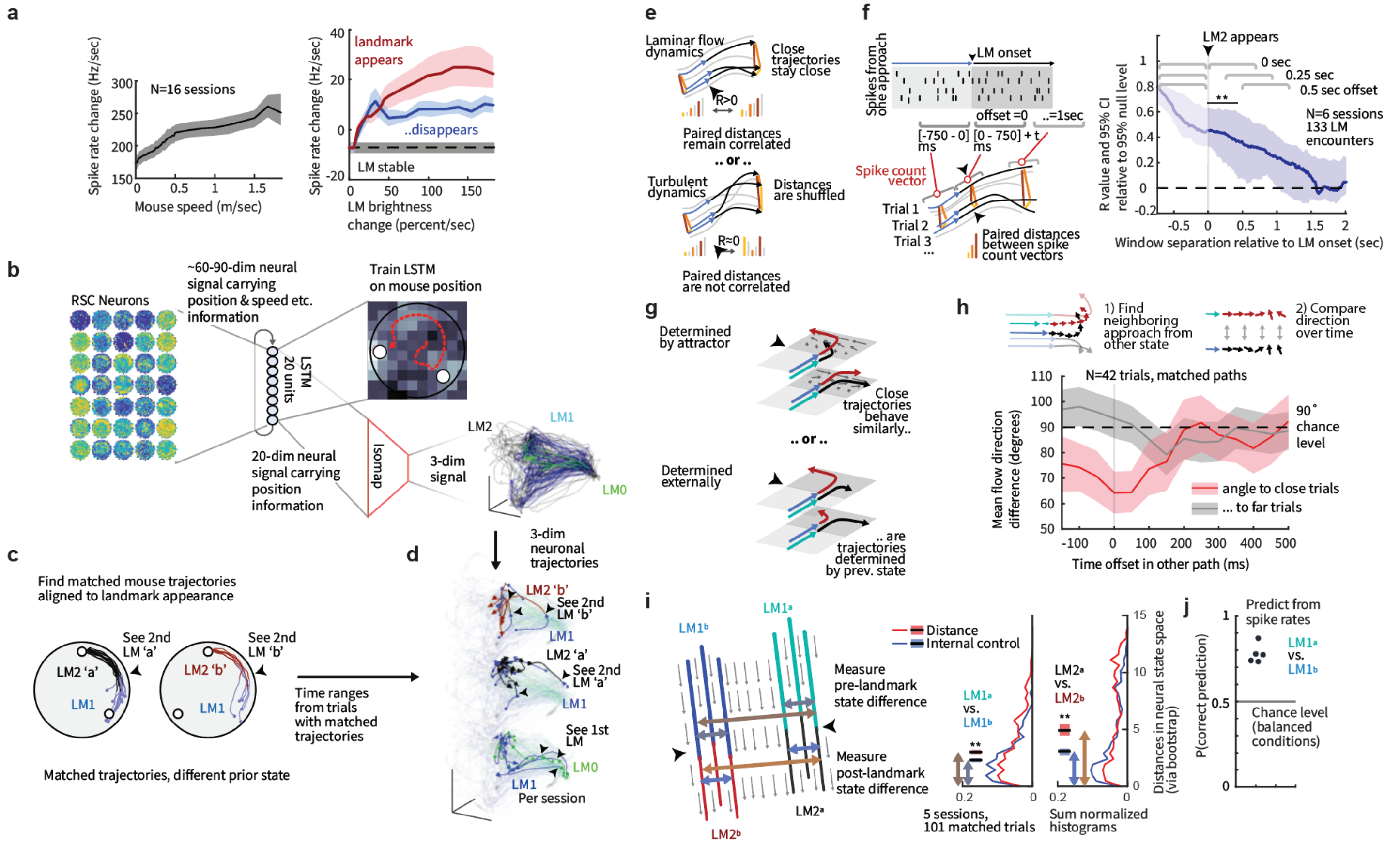


**Supplementary Figure 14.** Computation of low-dimensional spatial modes of mouse RSC activity. (**a**) Left: the speed at which the neural activity evolves (avg. speed of largest 5 principal components, filtered at 1Hz, CIs via bootstrap) correlates with running speed. Right: When landmarks appear/disappear, they perturb neural activity (effect of mouse speed is regressed out). (**b**) For some analyses of the low-dimensional dynamics in RSC (Fig. 4, Supplementary Fig. 13), rate fluctuations related to non-spatial covariates such as speed, heading, etc. were removed: a single-layer LSTM with 20 hidden units was trained to predict the mouse position in a 10x10 grid from the RSC rates. The network learned 20 spatially relevant mixtures of input firing rates, with appropriate temporal smoothing to represent the mouse location. These activations were then embedded into 3-D space via isomap[17]. (**c**) To find trials across which mouse trajectories as they approached the $2^{nd}$ landmark were similar, mouse trajectories were clustered (see Methods) leading to a subset of trials with similar locomotion and visual inputs. (**d**) The activity of RSC, in the low-dimensional representation, and in raw spike counts was then analyzed further. The example plot shows low-dimensional neural trajectories from LM0,1,2 states during matched mouse trajectories. (**e**) Alternative hypotheses for smoothness / predictability of neural dynamics across trials (corresponding to Fig. 4c). Dynamics across trials could behave like a laminar flow, so that trials with similar neural state remain so (top), or they could shuffle, leading to a loss of the pairwise distance relationships across trials (bottom). (**f**) We measured this maintenance vs. loss of correlation in a sliding 750ms window beginning at the $2^{nd}$ LM onset, versus a window just before. CIs were computed across sessions (See Methods). (**g**) Hypotheses for whether stable neural dynamics (Fig. 3b,c, Supplementary Fig. 13) can determine how RSC activity encodes disambiguated LM identity ('a' or 'b'). Top: trials in which the correct identity is 'a' but that are neurally close to other trials where the answer is 'b' might get dragged along in the wrong direction at least transiently. This would indicate relevance of recurrent dynamics on this computation. Bottom: alternatively, neural activity could be determined by the correct answer, even in trials that (in neural rate space) are close to trials from the opposing class. (**h**) We tested this by finding the closest trial from the opposing class (e.g. the closest $LM1_a$ for a $LM1_b$ trial) in RSC rate space. To evaluate co-evolution regardless of this selection confound, we then analyzed the direction of flow of the neural state over time (red). As a control, we also analyzed neurally far trials (grey). The flow direction of the neural activity was significantly aligned for ~100ms. (**i**) Left: Schematic for the analysis of representation of $LM1_a$ vs. $LM1_b$ states. Trial-to-trial distances were compared within group vs. across group. Right: Both before and after the $2^{nd}$ LM becomes visible, the classes are distinct in neural state space. (Same data as in Fig. 4d). (**J**) Whether a trial comes from $LM1_a$ or $_b$ can also be decoded from low-pass filtered (2Hz) firing rates before the $2^{nd}$ LM onset (via regression tree, cross-validated across trials, balanced N across conditions).

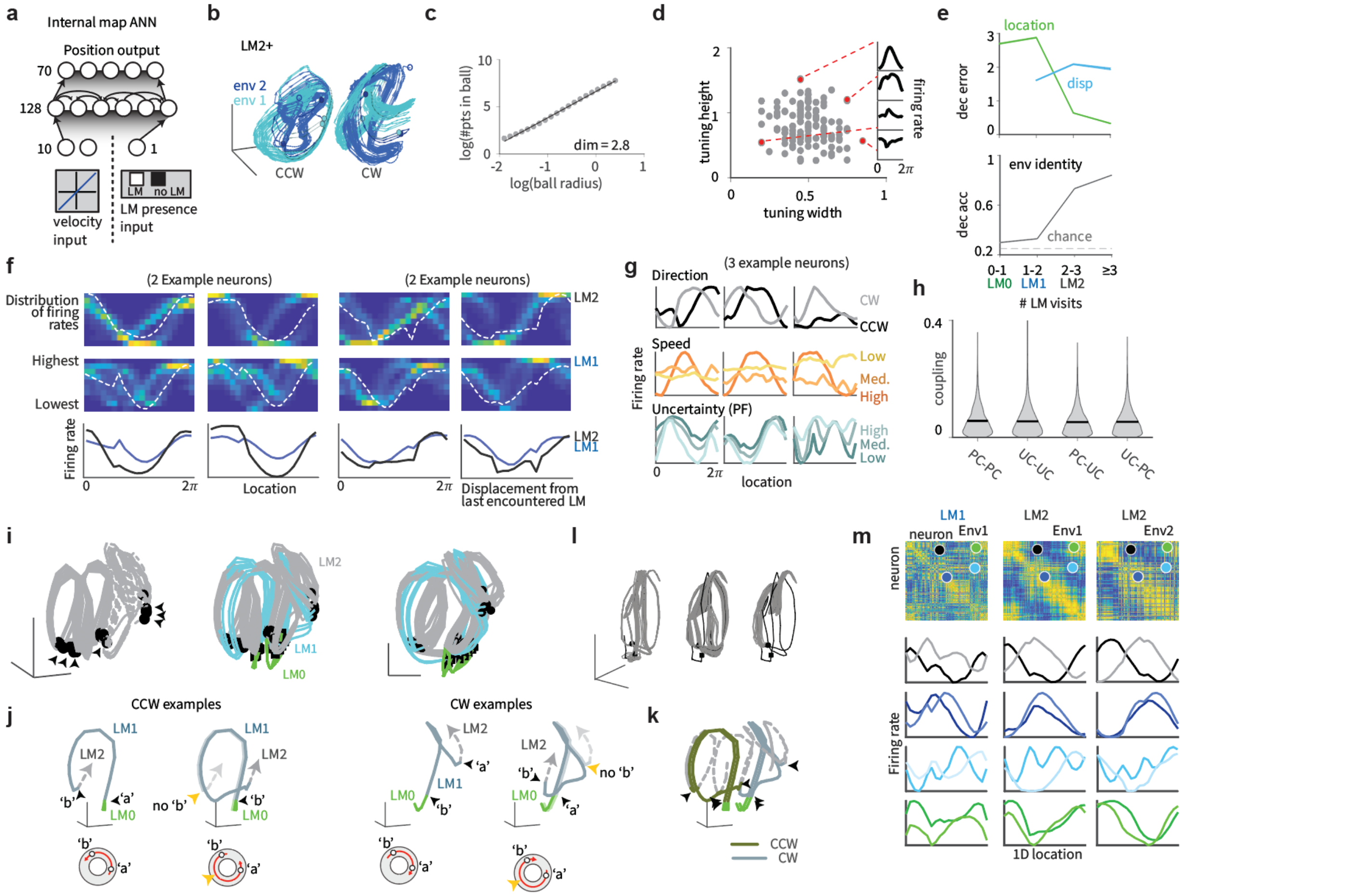

**Supplementary Figure 15.** ANN with binary landmark presence input recapitulates all main findings from the external map ANN. Here, the ANN must simultaneously infer the LM locations and the location of the animal, in contrast to the previous "external map" configuration. These determinations are inter-related, thus the much higher difficulty of the task. (**a**) Structure of the recurrent network. Input neurons encoded noisy velocity (10 neurons) and landmark (LM) information (1 neuron). In the internal map setup, the input signaled whether a LM was present at the current position or not. (**b**) State space trajectories in the internal map network after the second LM encounter in two different environments. The dark green / dark blue parts of the trajectories corresponds to the sections before the third LM encounter. Left: Predominantly counterclockwise trajectories, right: Predominantly clockwise trajectories. Landmarks and trajectories were sampled with the same parameters as experiment configuration 1, but the duration of test trials was extended from 10s (100 timesteps) to 50s (500 timesteps). Only trials with low error after the second landmark encounter are shown, defined as maximum network localization error smaller than 0.5rad, measured in a time window between 5 timesteps after the second landmark encounter until the end of the trial. Only the state-space trajectory after the second landmark encounter is displayed. (**c**) State space dimension is approximately 3, same analysis as in Supplementary Fig. 8c. (**d**) Example tuning curves, same analysis as in Supplementary Fig. 8b. (**e**) Linear decoding of position, displacement from last LM and LM separation from ANN activity, same analysis as in Supplementary Fig. 9c. A multinomial regression decoder was trained on 4000 trials from experiment configuration 1 (the training distribution of the internal map task) to predict from hidden layer activities which of the four possible environments was present. Performance was evaluated on separate 1000 test trials sampled from the training distribution. (**f**) Example neurons showing transition from egocentric LM-relative displacement coding to allocentric location encoding, same analysis as in Supplementary Fig. 9a,b. (**g**) Example neurons showing conjunctive encoding, same analysis as in Supplementary Fig. 5b. Location tuning curves were determined after the second landmark encounter using 1000 trials from experiment configuration 2 using 20 location bins. Velocity and uncertainty from the posterior circular variance of the enhanced particle filter were binned in three equal bins. (**h**) Distribution of absolute connection strength between and across location-sensitive "place cells" (PCs) and location-insensitive "unselective cells" (UCs), same analysis as in Supplementary Fig. 9e. (**i**) Hidden unit activations, corresponding to Fig. 2d. (**j**) Trajectories from example trials, same as in Fig. 2e. (**k**) Same trajectories as in j but with full LM2 state. (**l**) ANN is robust to perturbations, same as in Supplementary Fig. 12d. (**m**) ANN maintains pairwise correlation structure across states and environments, same as in Fig. 3a and Supplementary Fig. 8f.

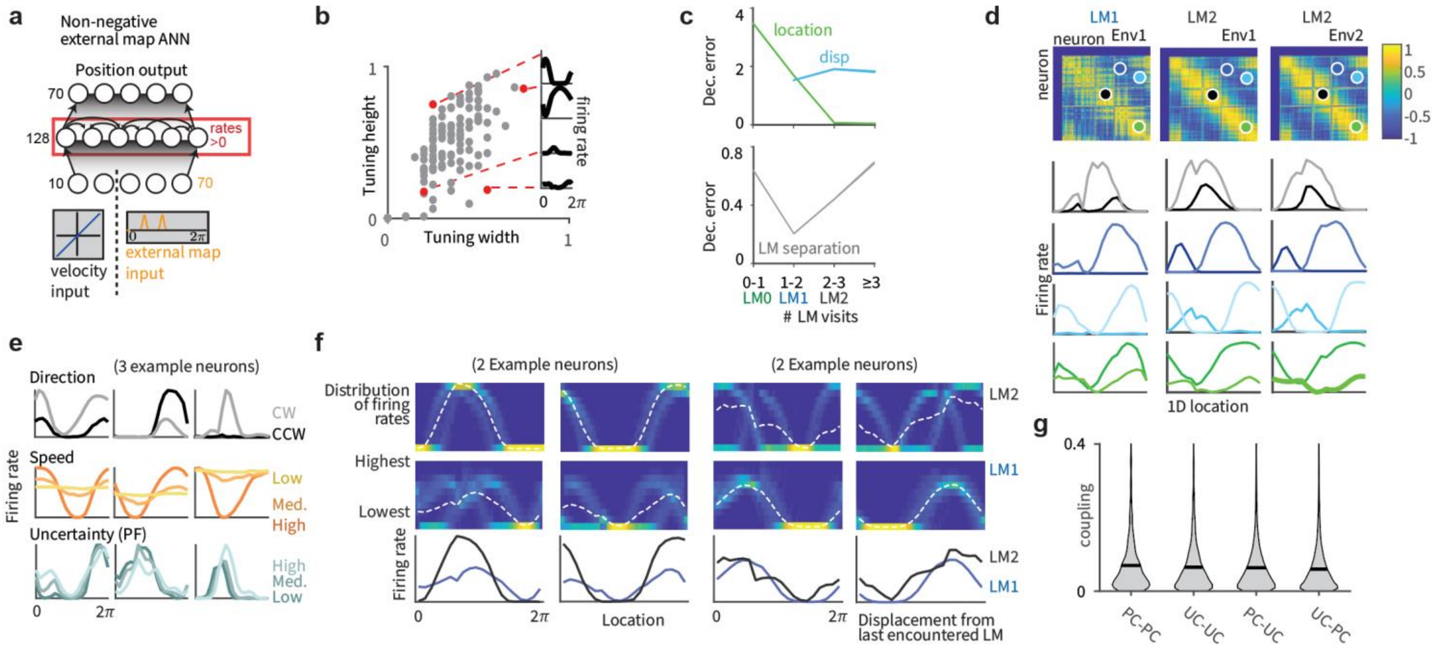


**Supplementary Figure 16.** ANN with non-negative rates recapitulates the main findings from the conventional ANNs. To probe the generality of the network predictions we imposed a non-negativity constraint on neural activity. Training an NN in the external map condition but with non-negative activity replicated all key results from the other NN types: we observed similar results with respect to location and displacement tuning (f), the transition in linear decodability of displacement to location from the population and dynamically varying decodability of LM separations within trials (c), the presence of heterogeneous and conjunctive tuning (e), lack of modularity in connectivity between cells with high and low amounts of spatial selectivity (g), and the preservation of cell-to-cell correlations across time within trials and across environments (d). The nonlinearity does affect the distribution of recurrent weights: The distribution of non-diagonal elements in the non-negative network is sparse (excess kurtosis k = 7.8), while it is close to Gaussian for the external and internal map networks with tanh-nonlinearity (k = 0.6 and k = 0.9 respectively; Supplementary Fig. 17a); however, the distributions of eigenvalues of the recurrent weights have similar characteristics for all trained networks (Supplementary Fig. 17b). (**a**) Structure of the recurrent network. Input neurons encoded noisy velocity (10 neurons) and received external map input (70 neuron), same as the regular external map ANN. Recurrent layer rates were constrained to be non-negative. (**b**) Example tuning curves, same analysis as in Supplementary Figs.15d. (**c**) Linear decoding of position, displacement from last LM and LM separation from ANN activity, same analysis as in Supplementary Fig. 15e. (**d**) ANN maintains pairwise correlation structure across states and environments, same as in Supplementary Figs. 8f and 15m. (**e**) Example neurons showing conjunctive encoding, same analysis as in Supplementary Figs. 5b and 15g. (**f**) Example neurons showing transition from egocentric LM-relative displacement coding to allocentric location encoding, same analysis as in Supplementary Figs. 9a,b and 15f. (**g**) Distribution of absolute connection strength between and across location-sensitive “place cells” (PCs) and location-insensitive “unselective cells” (UCs), same analysis as in Supplementary Figs. 8e and 15h.

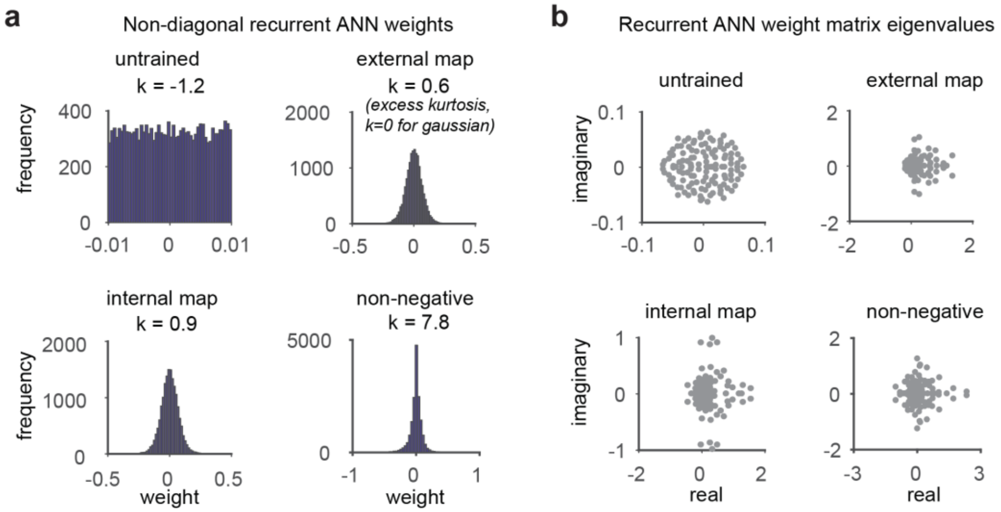


**Supplementary Figure 17.** ANN weights for the different architectures. The presence of a nonlinearity constraint on the ANN affects the distribution of recurrent weights: The distribution of non-diagonal elements in the non-negative network is sparse (excess kurtosis k = 7.8), while it is close to Gaussian for the external and internal map networks with tanh-nonlinearity (k = 0.6 and k = 0.9 respectively, panel a); however, the distributions of eigenvalues of the recurrent weights have similar characteristics for all trained networks (panel b). (**a**) Distribution of non-diagonal recurrent weights for randomly initialized (untrained), external map, internal map, and non-negative network. The k-value measured denotes excess kurtosis, a measure of deviation from Gaussianity (k = 0 for Gaussian distributions). (**b**) Scatterplot of real and imaginary part of complex eigenvalues of recurrent weight matrix for randomly initialized (untrained), external map, internal map, and non-negative network.

**Supplementary figures Bibliography:**


1. Voigts, J., Newman, J. P., Wilson, M. A. & Harnett, M. T. An easy-to-assemble, robust, and lightweight drive implant for chronic tetrode recordings in freely moving animals. *J. Neural Eng.* (2020) doi:10.1088/1741-2552/ab77f9.
2. Kropff, E., Carmichael, J. E., Moser, M.-B. & Moser, E. I. Speed cells in the medial entorhinal cortex. *Nature* **523**, 419–424 (2015).
3. Samsonovich, A. & McNaughton, B. L. Path Integration and Cognitive Mapping in a Continuous Attractor Neural Network Model. *J. Neurosci.* **17**, 5900–5920 (1997).
4. Burak, Y. & Fiete, I. R. Accurate Path Integration in Continuous Attractor Network Models of Grid Cells. *PLOS Comput. Biol.* **5**, e1000291 (2009).
5. Widloski, J. & Fiete, I. R. Inferring circuit mechanisms from sparse neural recording and global perturbation in grid cells. *eLife* **7**, e33503 (2018).
6. Welinder, P. E., Burak, Y. & Fiete, I. R. Grid cells: the position code, neural network models of activity, and the problem of learning. *Hippocampus* **18**, 1283–1300 (2008).
7. Jeewajee, A., Barry, C., O'Keefe, J. & Burgess, N. Grid cells and theta as oscillatory interference: Electrophysiological data from freely moving rats. *Hippocampus* **18**, 1175–1185 (2008).
8. Hardcastle, K., Ganguli, S. & Giocomo, L. M. Environmental Boundaries as an Error Correction Mechanism for Grid Cells. *Neuron* **86**, 827–839 (2015).
9. Ma, W. J., Beck, J. M., Latham, P. E. & Pouget, A. Bayesian inference with probabilistic population codes. *Nat. Neurosci.* **9**, 1432–1438 (2006).
10. Fiser, J., Berkes, P., Orbán, G. & Lengyel, M. Statistically optimal perception and learning: from behavior to neural representations. *Trends Cogn. Sci.* **14**, 119–130 (2010).
11. Zhang, K. Representation of spatial orientation by the intrinsic dynamics of the head-direction cell ensemble: a theory. *J. Neurosci.* **16**, 2112–2126 (1996).
12. Tsodyks, M. & Sejnowski, T. Associative memory and hippocampal place cells. *Int. J. Neural Syst.* **6**, 81–86 (1995).
13. Stefanovska, A., Strle, S. & Krošelj, P. On the overestimation of the correlation dimension. *Phys. Lett. A* **235**, 24–30 (1997).
14. Meshulam, L., Gauthier, J. L., Brody, C. D., Tank, D. W. & Bialek, W. Collective Behavior of Place and Non-place Neurons in the Hippocampal Network. *Neuron* **0**, (2017).
15. Yoon, K. *et al.* Specific evidence of low-dimensional continuous attractor dynamics in grid cells. *Nat. Neurosci.* **16**, 1077–1084 (2013).
16. Gothard, K. M., Skaggs, W. E. & McNaughton, B. L. Dynamics of Mismatch Correction in the Hippocampal Ensemble Code for Space: Interaction between Path Integration and Environmental Cues. *J. Neurosci.* **16**, 8027–8040 (1996).
17. Tenenbaum, J. B., de Silva, V. & Langford, J. C. A global geometric framework for nonlinear dimensionality reduction. *Science* **290**, 2319–2323 (2000).
